# Pre-existing Th1 immunity is abrogated by ongoing recruitment of monocytic host cells that are refractory to activation

**DOI:** 10.64898/2026.08.14.744955

**Authors:** Matheus B. Carneiro, Santiago Aguiar E. Soares, Chris Tiessen, Camila Gaio, Ben Perks, Leah S. Hohman, Bruna Araujo David, Paul Kubes, Matthias Mack, Ehud Inbar, Nathan C. Peters

## Abstract

Protective immunity against many infectious diseases develops following primary infection, called infection induced immunity (III), and provides a blueprint for vaccination. However, many vaccination strategies have failed. In the parasitic *Leishmania major* model of self-healing cutaneous disease, control of secondary challenge infection relies on pre-existing T helper (Th)1-dependent activation of skin-infiltrating monocytes for elimination of intracellular parasites. To better understand immune-evasion of pre-existing Th1 immunity by pathogens, we investigated the pathogen-niche established following non-healing challenge infection with the *L. amazonensis* parasite in a setting of pre-existing III. Following secondary challenge, pre-existing Th1 III initially controlled infection but ultimately failed. Loss of protection was not overtly STAT6- or IL-10-mediated. Rather, monocyte-lineage tracing revealed inflammatory monocyte-derived PD-L1^+^PD-L2^+^ macrophages provide an intracellular pathogen-niche and facilitate evasion of pre-existing Th1 immunity. Anti-PD-1 immune checkpoint blockade enhanced uninfected, but not infected, monocyte-derived cell activation and depletion of monocyte-derived precursors improved parasite control. These observations suggest that evasion of pre-existing Th1 immunity in this setting is not due to a failure of the Th1 response, but rather due to infected-cell intrinsic defects in activation.

## Introduction

Pathogens that cause chronic diseases have evolved mechanisms to establish infection and evade the subsequent generation of host immunity. The success of these evasion strategies potentially poses problems for efficacious vaccination as they may also mediate evasion of pre-existing immunity. This could explain the challenges associated with generating vaccines against many chronic infectious diseases. In the setting of chronic phagosomal pathogens, such as *Mycobacterium tuberculosis*, *Cryptococcus neoformans*, and *Leishmania spp*., these evasion strategies are associated with the establishment of intracellular pathogen niches within the phagolysosome of phagocytic cells(*1*). The definition of pathogen-niche adapted phagocytes *in-vivo* has been both elusive and dynamic with the increased appreciation of the myriad of phenotypes and functional plasticity that bone-marrow derived and tissue-resident phagocytes can adopt during infection(*2–4*). For example, different *Leishmania* species infect different phagocytic cell types, such as neutrophils(*5*), monocytes, monocyte-derived cells(*6–8*), embryonic derived macrophages(*9*), or multiple cell types(*10*). The definition of these phagocytic niches is further complicated by the fact that they are also associated with pathogen elimination given proper activation. In this context, CD4^+^ T helper 1 (Th1)-mediated immunity is essential to control self-healing *Leishmania* infections via Th1 cell-derived IFN-γ induced iNOS expression in infected mononuclear phagocytes(*11–14*), whereas IL-10 and the Th2 cytokines, IL-4 and IL-13, can inhibit this process via STAT-6 signaling and mediate Arginase I production(*15–17*). In settings of non-healing *Leishmaniasis,* determining if the parasite can establish a pathogen-niche even when robust pre-existing immunity exists and defining the phagocytic cells that provide this niche are critical factors in determining the nature and feasibility of efficacious vaccination or treatment. These insights are also likely relevant to other chronic phagosomal pathogens that employ phagosomes as sites of replication and persistence(*1*).

Primary infection with an infectious disease leading to the development of resistance to secondary exposure is known as infection induced immunity (III). This phenomenon has been described for infections by a diverse range of pathogens including viruses, bacteria, fungi, helminths and protozoan parasites(*18–27*). Infection-induced immunity provides the rationale for the development of vaccines, is a proof of their feasibility, and the specificity and class of immunity generated by protective III provides the blueprint for vaccine design. Thus, it is somewhat surprising that even though people develop III against many pathogens, there remains no efficacious human vaccines against numerous diseases that are associated with significant morbidity and mortality around the world, including tuberculosis, a large number of gastrointestinal infections, and numerous parasitic and vector-borne diseases, including malaria, schistosomiasis, and leishmaniasis (*28*). In the context of Leishmaniasis, the observation that III existed led to the practice of “Leishmanization” to induce immunity against the vector-borne intracellular protozoan parasite *Leishmania major.* This was achieved by inoculating naive individuals with live parasites from active cutaneous lesions without the disease enhancing features of vector-mediated transmission, leading to 90-100% protection against subsequent natural exposure(*29*). The protection associated with III against *Leishmania major* arises from concomitant immunity, where the chronic primary infection maintains a robust population of Ly6C^+^ Th1 effector (T_EFF_) cells that are rapidly recruited to the secondary challenge site and are sufficient to induce the recruitment and rapid activation of monocytes leading to parasite killing before establishment of the parasitic niche(*7, 28, 30, 31*). In some studies, CD4^+^ T resident memory cells can perform a similar function, although to a lesser degree(*32*).

In this work, we generated III employing a well-described self-healing model of cutaneous Leishmaniasis, in which C57BL/6 mice are inoculated sub-cutaneously in the footpad with 10^4^ *Leishmania major, L.m-*FV1, to mimic the practice of Leishmanization. This methodology generates superior Th1 protective immunity upon challenge versus intradermal inoculation or vaccine induced immunity (*27, 33–35*). We then assessed the capacity of this robust III to provide protection against challenge with non-healing species or isolates of *Leishmania* to determine how a phagosomal pathogen might evade pre-existing immunity. Employing an inducible lineage tracing model of CCR2^+^ monocytes, red fluorescent protein (RFP) reporter parasites to track infected cells, the *L. amazonensis* (*L.a.*) parasite, which is associated with a non-healing form of complicated cutaneous disease in South America(*36*), and back-to-back comparisons of *L. major* versus *L. amazonensis* sites of challenge infection, we were able to define a highly pro-inflammatory infected population of monocyte-derived macrophages that also expressed the immune checkpoint inhibitor ligands PD-L1 and PD-L2, the monocyte-recruiting chemokine CXCL9, and Arginase I. In contrast to uninfected monocyte-derived cells, these infected cells were refractory to immune-checkpoint blockade and provided a unique immune-evasive niche for *L. amazonensis* persistence and replication despite robust pre-existing Th1 immunity. Acquisition of this phenotype was dependent upon the time at which the monocytes were recruited into the tissue. Early arriving monocytes facilitated parasite control, but late arriving monocytes facilitated parasite expansion.

Our observations suggest that failure to control non-healing cutaneous Leishmaniasis following secondary challenge with *L. amazonensis* is not due to a defect in pre-existing Th1 immune response, but rather the refractory phenotype of pathogen-niche adapted monocyte-derived macrophages(*6, 37, 38*). This immune-evasive pathogen-niche may represent a critical target population for successful treatment strategies against phagosomal infections.

## Methods

### Ethics Statement

Ethics approval for this study was obtained from the Animal Care Committee (ACC) at the University of Calgary (Protocol number: AC23-0019) in compliance with the Canadian Council for Animal Care. Mice in this study were anesthetized using Ketamine(10mg/Kg) + Xylazine (1mg/Kg) prior to ear infection/challenge.

### Mice

C57BL/6J (Wt), *il10^-/-^* (B6.129P2-il10^tm1Cgn^/J), *stat6^-/-^* (B6.129S2C-stat6^tm1gGru^/J) and ZsGreen1 (B6.Cg-*Gt(*ROSA*)26Sor^tm6(CAG-ZsGreen1)Hze^*/ were originally obtained from The Jackson Laboratory and maintained under specific pathogen-free conditions at The University of Calgary Animal Resource Centre. The *il-10^-/-^stat6^-/-^* dKO mice were generated in house by crossing homozygous *il10^-/-^* and *stat6*^-/-^ mice. After genotyping the F2 generation and identification of homozygous *il-10^-/-^stat6^-/-^* mice by PCR, we maintained this colony by breeding homozygous dKO mice. The CCR2-creER^T2^ mice were kindly provided by Dr. Burkhard Becher (University of Zurich, CH) and the CCR2.CFP.DTR mice were kindly provided by Dr. Tobias Hohl (Memorial Sloan Kettering Institute, New York, NY). All mice were age and sex-matched and experiments were performed using mice between 5-12 weeks of age. Littermates of the same sex were randomly assigned to experimental groups. Mice were maintained at 5 animals per microisolator cages and *ad libidum* access to food and water.

### Parasites

Most experiments were performed using *L. amazonensis* (IFLA/BR/67/PH8) or the self-healing strain of *L. major* FV1 (MHOM/IL80/Friedlin) parasites. In addition, we performed specific experiments using the non-healing strains of *L. major* Ryan (WR2885) (*L.m_RYN*), originally isolated from human cutaneous lesion biopsies (*39*), and the non-healing *L. major* Seidman (MHOM/SN/74/SD) strain (*L.m_Sd*) isolated from a persistent human infection after treatment (*40*). A stable transfected line of self-healing *L. major* FV1 or *L. amazonensis* promastigotes expressing a red fluorescent protein (*L.m.-*RFP or *L*.*a.*-RFP) were employed and generated as described previously (*41*). Parasites were grown *in vitro* at 26°C in complete medium 199 (M199), supplemented as follow: 20% heat-inactivated FCS (Sigma-Aldrich), 100 U/ml penicillin, 100 μg/ml streptomycin, 2 mM L-glutamine, 40 mM Hepes, 0.1 mM adenine (in 50mM Hepes), 5 mg/ml hemin (in 50% triethanolamine), and 1 mg/ml 6-biotin. *L. amazonensis*-RFP was maintained in the presence of 50 μg/ml Geneticin (G418; GIBCO). Infectious metacyclic promastigotes were purified using a ficoll gradient and used for all injections (*42*).

### Generation of *Leishmania* hybrids

The parental strains of *Leishmania* parasites used to generate the experimental hybrids (2a, 4a and 6a) were *L. major* FV1 (MHOM/IL80/Friedlin) that has a blasticidin S-resistance (BSD) marker and *L. amazonensis* (IFLA/BR/67/PH8), that has a hygromycin B resistance marker. To generate the hybrids *Lutzomia longipalpis* sandflies were artificially feed through a chick skin membrane. An equal proportion of the parental strains of *Leshmania* parasites, with independent antibiotic resistance markers, were mixed in a heparinized mouse blood (4-8×10^6^ parasites in the promastigote logarithmic phase/ ml of blood). After 9 days the sandflies midguts were dissected, and hybrids parasites were recovered by double antibiotic selection, as previously described (*43, 44*). Three hybrids were used in this study based on their healing phenotype in the murine host.

### Primary Infection and Challenge

To generate III via the establishment of chronic primary infections, mice were injected with 10^4^ *L*. *major* metacyclic promastigotes subcutaneously in the hind footpad in a volume of 40μl and used at 12 to 20 weeks post-primary infection when footpad lesions had completely resolved. In the indicated experiments, chronic infected mice were also generated by injecting either 10^4^ *Leishmania* F1 hybrids *(L.m.xL.a): 2a, 4a 6a* or *with* 10^4^ *L. amazonensis* metacyclic promastigotes in the hind footpad in a volume of 40μl. Because primary *L.a.* infections in the footpad do not heal, animals with primary *L.a* infections were challenged at 8 weeks post-infection. Naïve mice and mice with a chronic primary infection, were challenged with either 2×10^5^ *L*. *major-*RFP, *L. amazonzensis*-RFP, *L. major*-RYN or *L.major*-Sd metacyclic promastigotes intra-dermally (i.d) in the ear in a volume of 10μl.

### Estimation of Parasitic Load at skin site

Parasite burden was determined by performing a two-fold serial dilution analysis in 96-well flat bottom plates containing M199. The number of viable parasites in each ear was determined from the highest dilution at which parasite growth was observed after 14 days of incubation at 26°C and results were expressed as the mean values of the negative logarithm of the titer.

### Processing of Tissue Ears

Ears were removed and placed in 70% ethanol for 5 minutes and then allowed to dry. The ventral and dorsal sheets of ears were separated and incubated in DMEM containing 160 μg/mL of Liberase (Roche Diagnostic) for 90 minutes at 37°C and 5% CO_2_. Digested ears sheets were homogenized for 40’ at 1714 rounds per run (rpr) using a gentleMACS Octo dissociator instrument (Miltenyi Biotec,Inc) with 6 ml DMEM media containing 0.05% DNase I and filtered using a 50 μm-pore-size cell strainer.

### Flow Cytometry

After tissue processing, cells were washed and labeled with Live/Dead fixable viability stain and anti-Fc III/II (CD16/32) receptor Ab (2.4G2) for 20 min at 4°C. This step was followed by a surface staining with different combinations of antibodies for 20 min at 4°C in the dark. For intracellular staining, cells were fixed with BD Cytofix/Cytoperm (BD Biosciences) and stained for 45 min at 4°C when staining cytokines and for 5 min at RT when only staining for the enzymes iNOS and Arg1. Data were collected from individual ears using a 5-laser Cytek Aurora (Cytek Biosciences) spectral flow cytometer and analyzed using FlowJo software (TreeStar). The follow anti-mouse antibodies were used: CD11b (M1/70), CD45 (30-F11); Ly6G (1A8); Ly6C (HK1.4); CD64 (X54-5/7.1); CCR2 (SA203611); CX3CR1 (SA011F11); CD11c (HL3); MHCII (M5/114.15.2); CD40 (3/23); CD206 (C068C2); iNOS (CXNFT); Arginase I (A1exF5); CD90.2 (53-2.1); TCR-B (H57-597); CD4 (RM4-5); CD8 (53-6.7); IFN-γ (XMG1.2); TNF-α (MP6-XT22); pro-IL1B (NJTEN3); PD-L2 (TY25); PD-L1 (MIH5); SiglecF (1RNM44N); CD24 (M1/69); CD49b (HMa2); CD200R3 (Ba13). All antibodies were obtained from Thermo Fisher, BD Biosciences or Biolegend. A portion of each sample was removed to determine the absolute number of cells using the AccuCheck Counting Beads (Thermo Fisher).

### Cytokine Analysis by Flow Cytometry

Single-cell suspensions obtained from skin were re-stimulated as previously described(*30*). Briefly, 10^6^/mL cells were stimulated with 0.5-1×10^6^ T cell-depleted (StemCell Technologies) naïve spleen cells (APCs) with 50 μg/mL freeze-thaw *Leishmania* antigen for 14 hours at 37°C in 5% CO_2_. During the last 7 hours of culture, 3 μg/mL of Brefeldin A (Sigma-Aldrich) was added. In some experiments, cells from the ear dermis were stimulated *ex-vivo* with PMA (Phorbol 12-myristate 13-acetate, Sigma Aldrich) 50 ng/mL and 1 μg/mL of ionomycin (Sigma Aldrich) for 4 hours. 10 μg/mL of Brefeldin A was added to the culture for the last 3 hours.

### Tetramer staining

PEPCK_335-351_:I-A^b^ tetramer staining was performed on single cell suspension after the tissue ear processing as previously described(*45*). Briefly, the skin samples were incubated for 1h at room temperature (RT) with either allophycocyanin (APC) or phycoerythrin (PE) labeled PEPCK_335-351_:I-A^b^ streptavidin tetramers.

### Real-time PCR

Ears cell homogenates from right and left ears were pooled and passed through QIAshredder columns. RNA was purified using a RNeasy min kit as per the manufacturer’s protocol (Qiagen). Reverse transcription was performed using the High-Capacity cDNA Reverse Transcriptional Kit (Thermo Fisher). Real-time PCR was performed on an ABI Prism 7900 sequence detection system (Applied Biosystems). The results were analyzed by the comparative threshold cycle method using 2^−ΔΔCT^ to determine the fold increase. Each gene was normalized to the 18S rRNA endogenous control and to non-infected mice as a sample control. TaqMan probes were used.

### Two-Photon Intravital Imaging

Image analysis was performed as described previously(*46*). Briefly, anesthetized mice were imaged in the lateral recumbent position, allowing the ventral side of the ear pinna to rest on a coverslip. Images were acquired using Leica SP8 two-photon microscope (Leica Microsystems), equipped with white light laser and resonance scanner and a 25X 0.95 NA water immersion objective. A combination of HyD-detectors and PMT internal (confocal) and external (2-photon) detectors were used to detect with 550-630 nm (RFP) and 490-550 nm (GFP). Captured images were processed using ImageJ 1.54g and Cellsens Dimensions® software. Images are single representative layers chosen from collected Z-stacks. RFP and ZsGreen1 channels were auto scaled to have equivalent maximal brightness. Contrast was optimized where appropriate to remove background.

### Tamoxifen Treatment

Tamoxifen (Toronto Research Chemicals) was dissolved in corn oil (Sigma) to 20 mg/ml and administrated in 100-150μl doses via intraperitoneal injections (100 mg/kg) as previously described(*6*) with two different schemes as explained in Fig 3.

**Fig 1.**
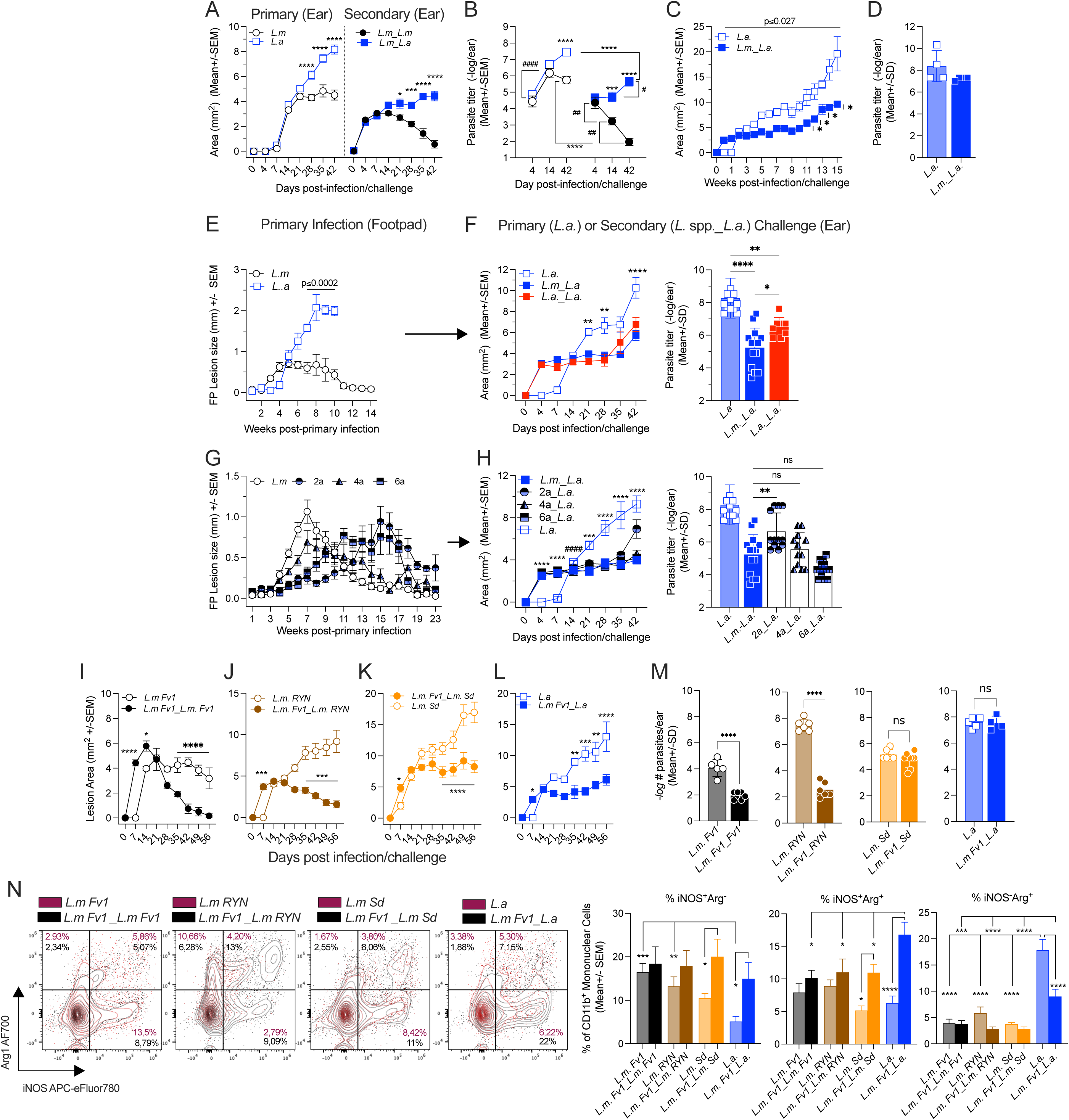
**Concomitant immunity provides short-term protection against heterologous challenge with *L.a***. (A-D) C57BL/6 mice were infected with 10^4^ self-healing *L. major* FV1 strain parasites in the footpad. After healing mice were challenged at a secondary intra-dermal (i.d.) site in the ear with 2×10^5^ *L. major* (*L.m._L.m.*) or *L. amazonensis* (*L.m_L.a.*) parasites. Age matched naïve control mice were also infected with parasites in the ear. Lesion area (A and C) and number (#) of parasites per ear determined by LDA (B and D) were assessed at the indicated time points post-infection/challenge (p.i./c). (E-F) For *L.a* homologous challenge (*L.a._L.a.*), C57BL/6 mice were infected with 10^4^ *L.a* parasites in the footpad (E) and 8 weeks later challenged i.d. in the ears with *L.a.* (F). (G-H) C57BL/6 mice were also infected with 10^4^ *L.m. X L.a.* F1 hybrids (2a, 4a and 6a) in the footpad (G). After healing mice were challenged with 2×10^5^ *L.a.* in the ears (H). Skin lesion area (F and H, left panels) and parasite load in the ear dermis at D42 p.c (F and H, right panels). (I-N) Infection Induced Immunity by *L. major*-FV1 provides variable levels of protection against *L. major* clinical isolates: C57BL/6 mice were infected with 10^4^ self-healing *L. major Fv1* in the footpad to establish III. After healing mice were challenged with either 2×10^5^ *L. major*-FV1 (I), *L. m*-RYN (J), *L.m*-Sd (K) or *L. amazonensis* (L) in the ears. Lesion areas were assessed over time (I-L) and # of parasites per ear (M) were determined at D56 p.i/c.. Phenotypical analysis of the % of Arg1 and/or iNOS mononuclear phagocytic cells at D56 p.i/c with the indicated parasite species was determined by FLOW (N). Data are from pooled experiments from 2 independent experiments (A-H) or from a representative of 2 independent experiments (I-N). n=3-5 mice per time point per experiment. *p≤0.01,**p≤0.0016, ***p≤0.0002, ****≤0.0001.

**Fig 2.**
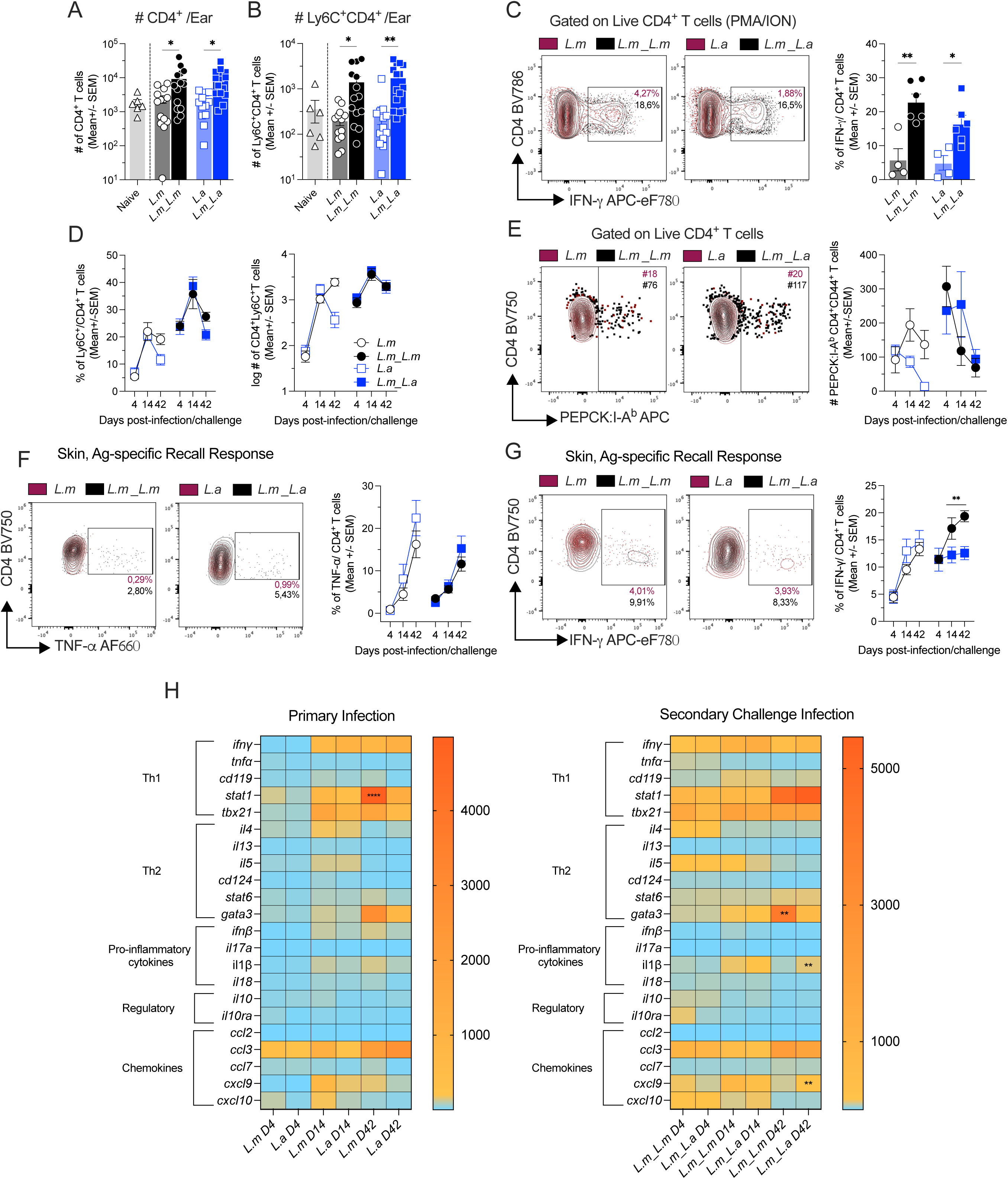
***L. amazonensis* challenge does not reduce the elicitation of Th1 immunity**. Primary and secondary infections were performed as described in Figure 1. Total numbers (#s) or frequency (%s) of total CD4^+^ T cells (A and D, left panel); Ly6C^+^CD4^+^T_EFF_ cells (B and D, right panel) or total #s of PEPCK:I-A^b^ CD4^+^CD44^+^ (E) cells per ear was determined employing quantitative flow cytometry. The % of total CD4^+^CD44^+^IFNγ^+^ cells was determined by FLOW after PMA+ION stimulation on D4 p.i/c. (C). Analysis of TNF-α (F) and IFN-γ (G) production by dermal CD4^+^ T cells following *ex-vivo Leishmania*-antigen re-stimulation including representative flow cytometry plots. Heat-map of expression levels of the indicated genes in the skin 4-, 14- and 42-days post-infection/challenge were determined by qRT-PCR for primary (H, left panel) or secondary infections (H, right panel). Data are from individual experiments (C-G) representative of ≥ 3 repeat experiments or pooled from ≥ 2 independent experiments (A, B and H). n=3-5 mice per time point per experiment. *p≤0.029, **p≤0.027, ****≤0.0001.

**Fig 3.**
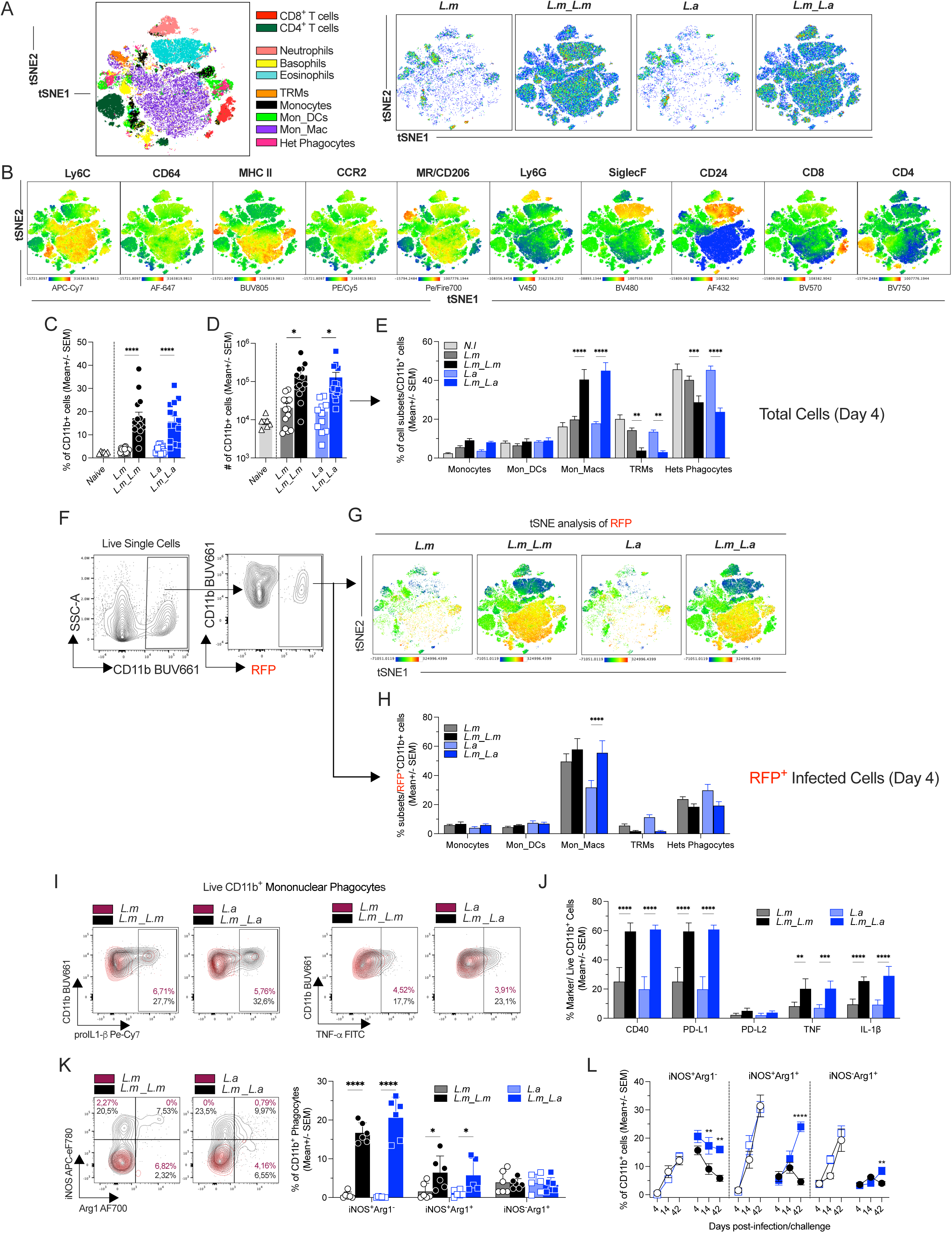
***L. amazonensis* heterologous versus *L. major* homologous challenge elicits an early equivalent immune response in settings of preexisting Infection Induced Immunity. tSNE map of skin immune cell clusters four days post-infection/challenge (p.i./c) with *L.m.* or *L.a.*** (A). tSNE map of overlay of indicated cell markers on immune cell clusters (B). % (C) and # (D) of total CD11b^+^ cells at the site of infection four days p.i./c as determined by flow cytometry. The % of the mononuclear phagocytic cell subsets within CD11b^+^ cells was determined by flow cytometry (E). Representative flow plots of CD11b and *Leishmania*-RFP (F) and tSNE map of RFP signal overlay of skin immune cell clusters (G). % of infected phagocytic cells was determined by tracking CD11b^+^RFP^+^ cells by FLOW (H). Phenotypic analysis of mononuclear phagocytic cells employing FLOW on d4 p.i/c (I-L). % of pro-IL-1β, TNF-α (I and J), CD40, PD-L1 and PD-L2 (J) and % of Arg1^+/-^ iNOS^+/-^ on d4 p.i/c (K) or on days −4,−14 and −42 p.i/c (L) in mononuclear phagocytic cells. Representative FLOW plots (I and K, left panels) of pro-IL1β, TNF-α and iNOS/Arg1. Data are from individual experiments (A, B, F, G) representative of ≥ 2 repeat experiments, or pooled from ≥ 2 independent experiments (C, D, E, H-L,). n=3-5 mice per experiment. *p≤0.02,**p≤0.01,***p≤0.0007, ****≤0.0001.

### Statistical Analysis

Results were expressed as mean ± SEM or SD as indicated. Student’s unpaired t-test were performed to determine differences between two groups. In cases where the variance between groups was different a Welch’s correction was employed. Comparisons between multiple groups were done using one or two-way ANOVA with Sidek’s post-test to correct for multiple comparisons. For data that was not normally distributed, data was compared employing a Kruskal-Wallis test with Dunn’s post-test to correct for multiple comparisons. Absolute number data were log transformed before statistical analysis. Data from repeat experiments were pooled when possible. Exact value of n, precision measures (mean + SD or SEM) and statistical significance are reported in the Figures and Figure Legends. All analyses were conducted using GraphPad Prism version 10.

### Monocyte depletion

Monocytes were depleted in two different ways. As previously described(*7*), we administrated Diphtheria toxin (DT) to CCR2-DTR mice during chronic time point of either primary or secondary infection with *L.a* parasites, as indicated in Fig 5. (DT) was obtained from Millipore, reconstituted at 1 mg/ml in PBS, and frozen at −80°C. Mice received 20 ng/g DT via the intra-peritoneal route in 0.2 ml PBS. The toxin was injected every two days, in a total of five doses, to CCR2-DTR mice during chronic phase of *L.a* infection/challenge. Littermate controls were used and given DT. All mice were euthanized 48h after last DT treatment. Monocytes were also depleted by using the anti-CCR2 antibody (Clone: MC-21(*47*)) kindly provided by Dr. Matthias Mack. Monocytes were depleted by daily injection of 20ug of MC-21 i.p. for five consecutive days and mice were euthanized 48h after the last injection. Control mice received administration of daily 20ug of rat IgG2b kappa (BioXCell) isotype control.

**Fig 4.**
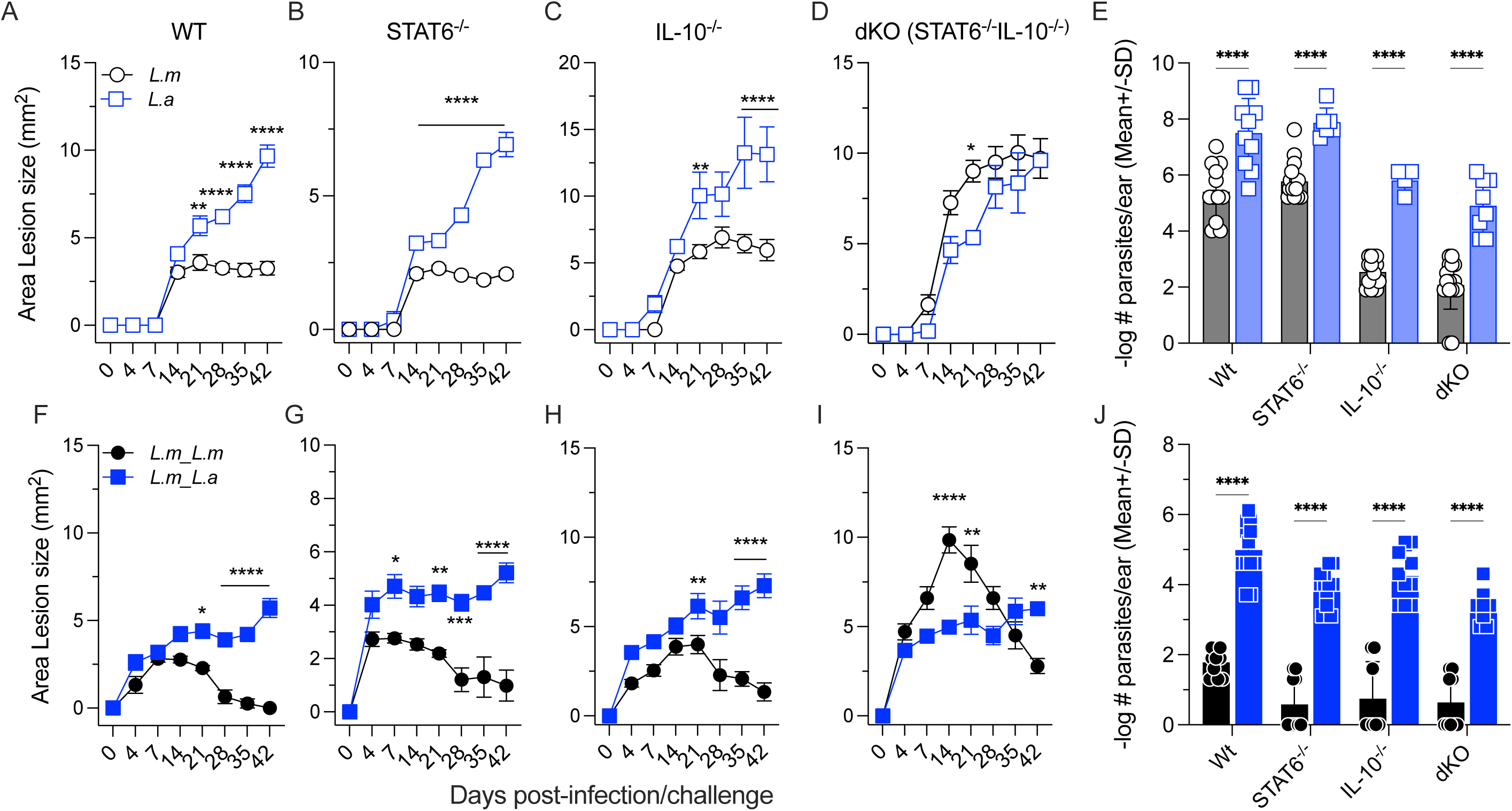
Absence of STAT6 signalling and/or IL-10 does not lead to enhanced protection after *L.a.* challenge. Primary and secondary infections were performed as described in Figure 1 employing C57BL/6 Wt or *stat6^-/-^, il10^-/-^* or *stat6^-/-^il10^-/-^* (dKO) mice. Lesion area (A-D, F-I) and # of parasites per ear (LDA) (E, J) were determined at D42 after *Leishmania spp.* primary or secondary infection. Data are from 2 independent pooled experiments. n=3-5 mice per experiment. *p≤0.04,**p≤0.0017,***p≤0.0002, ****≤0.0001.

**Fig 5.**
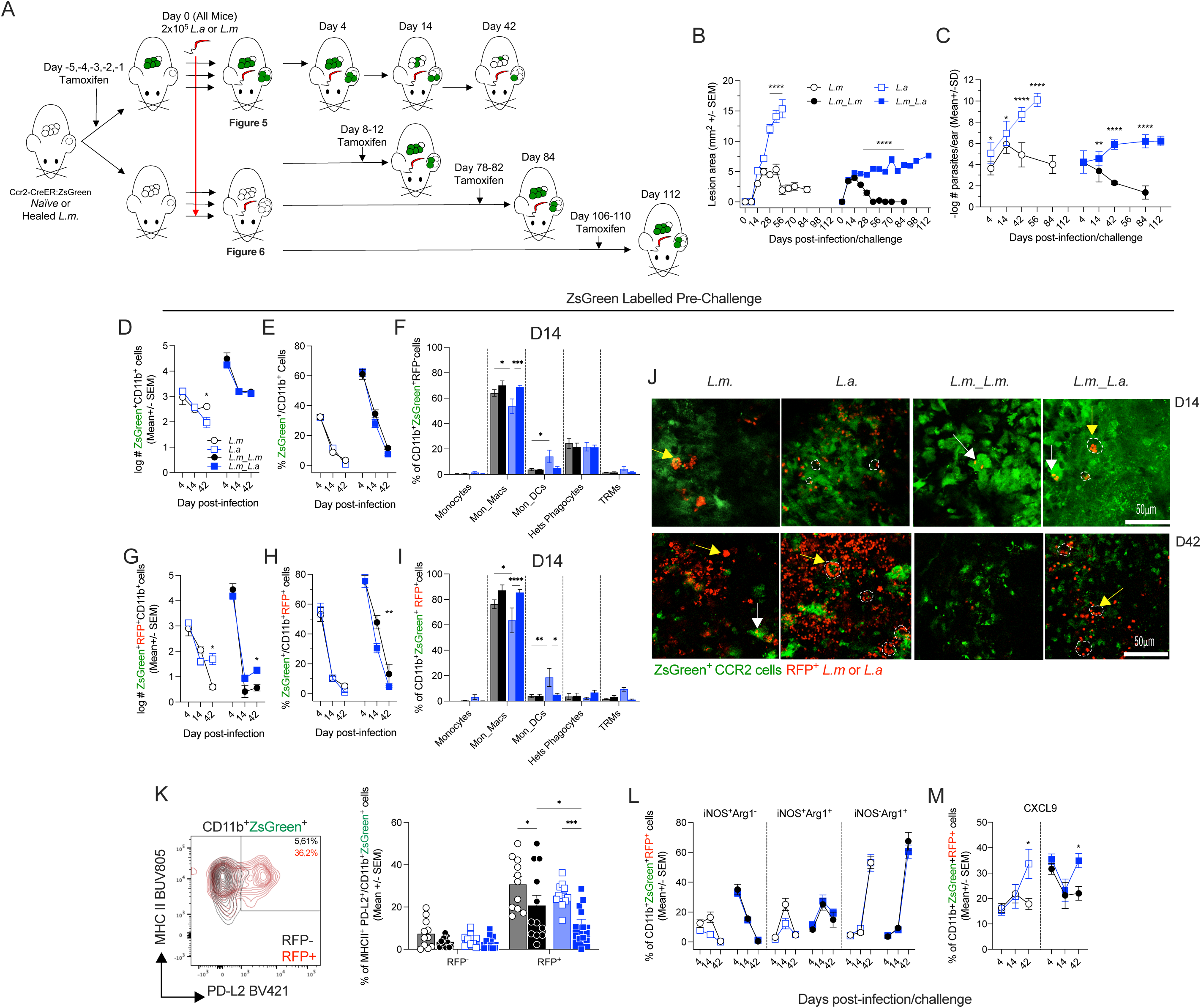
Monocyte fate tracing pre-*Leishmania* challenge. Schematic representations of tamoxifen (Tmx) treatment, pre or post-challenge and analysis of CCR2-creER^T2^.ZsGreen1 mice (A). Lesion area over-time (B) and # of parasites (C) was determined at the indicated time points. The total #s (D) or % (E) of CD11b^+^ZsGreen^+^ cells and the %s of the indicated cell subsets were determined by FLOW (F). Total #s of infected RFP^+^CD11b^+^ZsGreen^+^ cells per ear (G) and % of RFP^+^CD11b^+^ (H) that were ZsGreen^+^ at the indicated time points. % of indicated cell subsets within total CD11b^+^ZsGreen^+^RFP^+^ cells 14 days post-challenge (I). Visualization of ZsGreen^+^ cells at dermal site of *L.m*-RFP^+^ or *L.a*-RFP^+^ infection employing 2P-IVM at the indicated time points (J). The % of MHC II and PD-L2 within the RFP^+^ or RFP^-^ ZsGreen^+^CD11b^+^ population was determined by FLOW at D14 post-challenge (K); and the % of Arg1 and/or iNOS cells (L) or the % of CXCL9^+^ (M) within the RFP^+^ZsGreen^+^CD11b^+^ population was determined by FLOW at indicated time points. Data are from pooled experiments from ≥ 2 independent experiments (A-I and K-M), or from individual experiments (J) representative of ≥ 2 repeat experiments. n=3-5 mice per time point per experiment. *p≤0.03,**p≤0.0095,***p≤0.001, ****≤0.0001.

### *In vivo* anti PD-1 treatment

Fourteen days after the infection/challenge with *L.amazonensis*, C57BL/6 mice were given 150 μg of anti-PD-1 (clone RMP1-14, BioxCell) or the isotype control (rat IgG2a isotype control anti-trinitrophenol, clone 2A3). Mice were treated twice a week for 4 weeks, for a total of 8 doses, and euthanized at d42, two days after the last dose. A different antibody clone for PD-1 was used for the FLOW cytometry analysis (clone 29F.1A12).

## Results

### *L.a.* secondary challenge abrogates pre-existing Th1 immunity mediated by prior infection

We employed a validated model to establish pre-existing Th1 III in which C57BL/6 mice were inoculated in the footpad with a self-healing strain of *L. major* (*L.m*_FV1) resulting in a healed but chronic primary infection (Fig S1A and Fig S1B). Mice with healed primary infections are highly resistant to homologous secondary challenge with *L.m.*-FV1(*27*) and in many cases heterologous secondary challenge (Ref.(*48*) and references therein). To evaluate the protective capacity of III against secondary challenge with a strain of *Leishmania* that causes non-healing cutaneous disease, healed mice underwent heterologous challenge at a distal dermal site in the ear with *L. amazonensis* (*L.a)* parasites (*L.m.*_*L.a.*), or homologous *L.m* parasites as a positive control (*L.m.*_*L.m.*). We employed a challenge dose of 2×10^5^ via the intra-dermal route as this dose, while higher than that deposited by vector-mediated transmission (1-10 000)(*49*), more accurately recapitulates the inflammatory response observed following sand fly mediated transmission in a more tractable experimental model(*50*). This dose also allows for reliable tracking of RFP^+^ infected cells by flow cytometry, a critical component of our analysis. Following an early equivalent phase of lesion progression comparing *L.m._L.a.* (closed blue squares) versus *L.m._L.m.* (closed black circles) on days 0-14 (Fig. 1A), lesions in *L.m._L.a.* mice ultimately did not heal, and parasite numbers actually increased between days 14 and 42 (Figs 1B), resulting in an approximately 5000X increase in parasite load per lesion versus *L.m._L.m.* mice. In *L.m._L.m.* mice, lesions fully healed (filled black circles, Fig. 1A) and parasite load steadily decreased over the course of the experiment (Fig. 1B). Therefore, despite some degree of protection compared to mice undergoing a primary *L.a.* infection (open blue squares, Fig. 1, A and B), at no point did III reduce the lesion size or parasite burden following *L.a.* challenge. To determine if III simply needs more time to clear *L.a.* parasites we tracked *L.a.* challenge infections for 15 wks. At approximately 9 weeks, lesion growth returned to a progressive uncontrolled phenotype, significantly increasing week over week (Fig 1C). At 15 wks p.i., an equivalent number of parasites were found in the skin of *L.m._L.a.* mice compared to *L.a* primary infection (Fig 1D), demonstrating a complete loss of protection. Our observations are similar to previous studies in which III slowed progressive disease but did not mediate healing at *L.a.* secondary challenge sites and/or result in an appreciable reduction in parasite numbers over time(*51, 52*). We have now followed the course of challenge infection for a longer period to demonstrate that, after the initial period of slowed progressive disease, protection is completely lost after 12 weeks post-challenge and returns to a fully progressive phenotype with equivalent parasite loads. Therefore, slowing progressive disease does not ultimately lead to protection.

To more carefully determine if III generated by *L.m.-*FV1 represents the most protective infection induced immunity model available we also employed homologous *L.a.* or *L.m.* (FV1) x *L.a.* (PH8) F1 hybrid parasites to induce III. In settings of concomitant immunity, progressive disease at a primary site of infection can co-exist with control at a site of secondary challenge(*53, 54*) (see also Fig. S1, C and D). Therefore, we performed homologous challenge after generating III with *L.a* via primary inoculation in the footpad, which induced a non-healing but stable primary infection (Figure 1E). However, III induced by *L.a.* did not lead to improved protection against *L.a.* secondary challenge versus that mediated by *L.m.,* and, in fact, at 6 weeks post-challenge these mice had higher parasite loads (Fig 1F). This data suggests that simply priming the T cell response with a homologous antigenic repertoire is not sufficient to improve upon the III mediated by *L.m..* To investigate this further, we also generated *L.m.* (FV1) x *L.a.* (PH8) F1 hybrid parasites to induce III. Of the hybrids generated, we employed three that achieved a healed primary infection in the footpad (Fig 1G), similar to the *L. major* parental line. Based on the inheritance characteristics of *Leishmania,* these hybrids potentially possess a full complement of both *L.m.* and *L.a.* antigens(*55*). None of the hybrids improved upon the III generated by the parental *L.m.* strain following *L.a.* challenge in the ears, and the 2a strain was worse (Fig 1H). These data demonstrate that III mediated by *L.m.* infection provides protection that is on par or superior to that provided by homologous *L.a.* or hybrid parasites. They also suggest, but do not definitively prove, that an overt deficiency in unique *L.a.* antigens is unlikely to be the limiting factor in the failure of *L.m.*-mediated III.

### Infection Induced Immunity by *L. major*-FV1 provides variable levels of protection against *L. major* clinical isolates

We next wished to extend our observations and determine if the inability of *L.m.* FV1-mediated III to provide protection was unique to *L.a.* challenge. We challenged healed C57BL/6 mice with two *L.m.* clinical isolates known to produce more severe cutaneous disease, *L.m.*-RYN (*34, 39, 56*) and *L.m.*-Sd (*54*). Despite causing progressive disease with extremely high parasite loads in naïve mice, mice with III challenged with *L.m*-RYN parasites were able to control the disease as well as the homologous challenge with self-healing *L.m*-FV1 parasites, as seen by healing lesions (Figs 1, I and J), robust parasite control (Fig 1M), and a profile of iNOS and Arginase production by phagocytes that resembled *L.m.*-FV1 challenge sites (Fig. 1N). Protection against *L.m.*-RYN represents the ideal scenario for live or attenuated vaccine platforms, where III mediates protection against a highly virulent clinical isolate. On the other hand, mice challenged with *L.m*-Sd parasites did not resolve the infection, as seen by the same steady lesion size over time (Fig 1K), similar to *L.a* challenged mice (Fig 1L), and equivalent parasite numbers (Fig 1M). The lack of control of *L.m.*-Sd occurred despite the fact that in naïve mice this parasite did not cause excessive parasite loads (Fig 1M) and was associated with a profile of iNOS and Arginase producing cells (Fig 1N) similar to that of FV1 and RYN challenged mice, suggesting that the parasite has evolved mechanisms to evade Th1 immunity as suggested by previous studies of its preference for tissue-resident macrophages (TRMs)(*57*), see Discussion. Of interest, *L.a.* challenge was associated with greater frequencies of Arg1^+^ phagocytes, which did not occur with the *L.m.* strains even under conditions of similar parasite loads, suggesting a unique association with *L.a.* infection. Given the importance of understanding the mechanisms by which pathogens evade pre-existing immunity to formulating efficacious vaccines and the high clinical burden of *L.a.* infection in infected individuals, we continued our investigation employing *L.a.* challenge.

### Expression of early CD4^+^ Th1 infection induced immunity is intact after heterologous (*L.m_L.a*) secondary challenge

Rapid (≤4 days) recruitment and IFN-γ production from pre-existing Ly6C^+^ T_EFF_ cells is critical for protection mediated by pre-existing Th1 III at dermal and visceral sites of *Leishmania* homologous and heterologous secondary challenge(*30, 58*). We have argued previously that day 4 is a defining timepoint for analysis(*7, 30, 31, 59*). We hypothesized that the eventual loss of protection seen following *L.a.* challenge infection is due to a defect in elicitation of early CD4^+^ T cell mediated immunity in this setting. However, analysis of the CD4^+^ T cell response revealed higher #s of total CD4^+^ T cells (Fig. 2A), Ly6C^+^ expressing CD4^+^T_EFF_ cells (Fig 2B), and IFN-γ production (Fig. 2C) at sites of *L.a.* challenge versus primary infection, and these responses were equivalent to those at sites of *L.m.* secondary challenge. Over time, the frequency and number of Ly6C^+^CD4^+^ T cells remained equivalent at *L.m.* versus *L.a.* secondary challenge sites (Fig. 2D). We extended our findings by tracking *Leishmania*-specific CD44^+^PEPCK:I-A^b^ tetramer^+^ cells(*60*) (Fig 2E) and polyclonal cells producing TNF-α (Fig 2F) and IFN-γ (Fig 2G) after *ex-vivo Leishmania*-antigen specific stimulation. No overt differences in the number of tetramer^+^ cells or capacity of CD4^+^ T cells to produce IFN-γ or TNF-α in response to stimulation was observed. Interestingly, while the early day 4 IFN-γ response was equivalent in *L.a.* versus *L.m.*-challenged mice (Fig. 2G, right panel) and greater than that observed in primary infections, responses were significantly lower on days 14 and 42, despite equivalent or higher parasite loads (Fig. 1B). We also performed a real-time qPCR analysis of the skin on days 4, 14 and 42 post-challenge (Fig 2H). Both homologous and heterologous secondary challenge groups also showed increased expression levels for genes associated with Th1 (*ifnγ*, *stat1* and *tbx21*) Th2 (*il4*, *il5*, *stat6* and *gata3*) and chemotaxis response (*ccl3*, *cxcl9* and *cxcl10*) versus primary infection at D4 (Fig 2H, right panel, p≤0.05). Surprisingly, despite having bigger lesions and more parasites at the site of infection, mice challenged with *L.a*, showed remarkably similar levels of expression of key genes 14 and 42 days after infection, such as *ifnγ, tnfα, tbx21, il4, il10, stat1, stat6, cxcl10* compared to *L.m._L.m.* mice. Amongst the 22 genes analyzed only *il1β* and *cxcl9* were upregulated in the *L.m._L.a.* versus *L.m._L.m.* group on day 42 post-challenge, while the *L.m._L.m.* group induced higher levels of *gata3* at day 42 post-challenge. While our data indicates that the acute expression of III following heterologous *L.a.* challenge is largely intact, the equivalent expression of Th1 associated cytokines and genes despite equivalent or higher parasite loads at later time points suggests a deficiency in the elicitation of CD4^+^ Th1-mediated immunity at later time points may be occurring.

### Early monocyte and monocyte-derived phagocyte activation is intact after heterologous (*L.m_L.a*) secondary challenge

Early recruitment and activation of monocytes and monocyte-derived cells is critical for protection mediated by III at dermal sites of *L.m._L.m.* challenge (*7, 30, 31, 59*). To determine if *L.a.* challenge was associated with a deficit in phagocyte activation we conducted a thorough analysis of the phagocyte population on day 4 post-challenge. We employed a t-Distributed Stochastic Neighbor Embedding (tSNE) analysis of our spectral-flow panel employing fifteen key phenotypic markers (CD11b, CD45, Ly6C, CCR2, Ly6G, CD64, MHC II, CD206, CD24, SiglecF, CD8, CD4, TCR-β, CD49b, CD200R3) to identify 10 clusters of cells as defined in Fig S2A (*6, 9, 61*) and shown as a 2D scatterplot in Fig 3A (left panel). A comparison of the different challenge conditions employing tSNE analysis is shown in Fig. 3A, right panels). Expression of 10 specific lineage-defining markers on each cell subsets is shown in Figure 3B (Compare the expression of a single marker in 3B with the corresponding cell type defined in 3A). We observed a robust increase in the frequency and number of total myeloid CD11b^+^ cells at sites of secondary versus primary challenge (Fig 3, C and D). However, no differences were observed in the frequency, number, or composition of the innate CD11b^+^ immune cell infiltrate when comparing *L.m.*_*L.a.* versus *L.m.*-*L.m.* (Fig. 3. C-E) at day 4 post-challenge. Sites of secondary challenge were defined by an increase in the frequency (%) of monocyte-derived macrophages (Mon_Macs). These are defined as CD64^+^Ly6C^lo/hi^CCR2^-^ cells, do not include tissue-resident macrophages (TRM)^-^, but do include the P4-P5 monocyte-derived macrophage populations referred to by others(*61*). Increased Mon_Mac frequencies were associated with a corresponding decrease in the % of mature resident cell populations, such as mannose receptor (MR)^+^ embryonic derived TRMs and a less characterized population of CD11b^+^ phagocytic cells (CD64^-^ Ly6C^-^), onwards referred to as Heterogenous phagocytes (Hets_Phagocytes). These Hets_Phagocytes can have either a pre-existing tissue resident or monocyte derived origin depending upon steady-state or inflammatory conditions (*6, 61*) (Fig 3E). Numerically, while the numbers of TRMs did not change, more monocyte-derived-cells (Mon_Macs and Het_phagocytes) and recently recruited monocytes (CD64^-^ Ly6C^hi^CCR2^+^) were observed at sites of secondary challenge (Fig S2C). In all cases, we did not observe any significant differences between homologous (*L.m._L.m.)* and heterologous (*L.m._L.a.)* challenged mice at day 4 post-challenge.

Using *L.m-*RFP^+^ and *L.a*-RFP^+^ parasites to track the phenotype of infected cells at D4 post-challenge (Fig 3F) we also found no differences in the infected cell phenotype in *L.m._L.a.* versus *L.m._L.m.* challenged mice, and most infected cells for all conditions were Mon_Macs (Fig 3G and 3H). Associated with the early influx of IFN-γ-producing T cells (Fig. 2), we also observed that CD11b^+^ mononuclear phagocytic cells at homologous and heterologous secondary challenge sites bore markers of IFN-γ exposure including increased % of CD40, PD-L1 and TNF-α and pro-IL1-β but not PD-L2 (Figs 3, I and J, and Fig S2B, for dot plots). Lastly, the % of iNOS^+^Arg1^-^ or iNOS^+^Arg1^+^ mononuclear phagocytic cells was also increased in both secondary challenged groups (Fig 3K). These observations demonstrate that, similar to homologous challenge with *L.m.*, heterologous challenge with *L.a* parasites induced robust skin infiltration, activation, and maturation of monocytes on day 4 post-challenge, including increased expression of markers such as CD40 and iNOS, molecules important for parasite control(*62*). We also assessed the frequency of iNOS and Arginase producing phagocytes over time and found that secondary *L.a.* challenge induced significant frequencies of iNOS^+^Arg^-^ cells on day 14 and 42, consummate with parasite loads (Fig. 1B), but this was accompanied by increased Arginase production in the total phagocyte population (either iNOS^+^Arg^+^ or iNOS^-^Arg^+^) (Fig. 3L) by day 42 post-challenge, similar to our observations in Fig. 1N. Phagocyte arginase production has been associated with parasite persistence and proliferation(*63, 64*). Taken together, and similar to our analysis of the CD4^+^ Th1 response in Figure 2, protective immunity appears be intact at day 4 post *L.a.* heterologous challenge. However, at later time points, and in particular at day 42, the response begins to show signs of divergence from the ultimately protective response observed at sites of *L.m.*_*L.m.* challenge.

### Simultaneous abrogation of IL-10 and STAT6 signaling has a minor impact on disease control after *L.a.* challenge

During the later stages of infection in *L.m_L.a* challenged mice, the % of phagocytes expressing Arginase increased (Fig. 3L), despite a robust Th1 environment (Figs 2 and 3). Regulation of phagocytic cells by IL-10 and Th2 associated cytokine (IL-4 and IL-13)-mediated STAT6 signaling can induce an arginase-positive phagocyte phenotype and prevent intracellular pathogen killing (*6, 65, 66*). We wished to investigate a potential role for IL-10 and Th2 cytokine mediated STAT6 signaling in the lack of long-term protection following *L.a.* challenge and determine if removing these signaling pathways would result in a level of protection more analogous to that observed in *L.m._L.m.* mice. We established III in C57BL/6 Wt, *stat6^-^*^/-^, *il10*^-/-^ and dKO (*stat6^-/-^il10^-/-^*) mice via inoculation of the *L.m*-FV1 strain in the footpad. After homologous challenge with *L.m.,* the lack of signaling mediated by IL-10, STAT6, or both (dKO), did not alter the healing phenotype (compare open and closed circles in Fig 4, A-D and F-I), and led to a further reduction in the already low parasite loads, reaching undetectable numbers in a significant number of mice (Fig 4E and 4J). After secondary *L.a* challenge IL-10 and/or STAT6 signaling did not reverse the non-healing phenotype (closed squares in Fig 4, F-I), with a trend towards progressive disease at later timepoints, failed to reduce parasite loads to those levels observed following secondary *L.m.* challenge of KO or WT mice (Fig. 4J), and did not improve upon the fold-change reduction observed in *L.m.*_*L.a.* versus *L.a.* WT mice (Compare squares between Fig. 4, E and J). Collectively, our data indicate a significant, but minor role played by IL-10 and STAT-6 in mediating the loss of protection during *L.a.* heterologous challenge.

### CCR2-Lineage-Tracing reveals infected monocyte-derived phagocytes recruited at the time of *L.a.* challenge maintain a protective phenotype

Because monocytes undergo significant maturation once arriving at the dermal site of infection and loose expression of defining markers such as CCR2 and Ly6C, their fate cannot be easily defined over time (*17*). Therefore, we employed a tamoxifen (Tmx)-inducible lineage tracing model (CCR2-creER^T2^:Ai6(RCL-ZsGreen1 mice) in which Tmx treatment permanently turns on ZsGreen fluorescent protein expression based on CCR2-expression prior to maturation, predominantly in the blood. To track the fate of the initial wave of recruited monocytes, mice were treated with Tmx just prior to challenge (Fig. 5A, top row) or, to track the fate of monocytes recruited into the skin at later timepoints, mice were treated with Tmx at three time points after challenge (Fig 5A, bottom row, and Fig. 6) and lesion size and parasite loads were assessed (Fig 5, B and C). Note that mice in the *L.a.* group were euthanized at day 56 because they reached their humane endpoint. Quantification of the total number of phagocytes and phagocyte subsets per ear over the course of the entire experiment can be found in Figure S3A and S3B.

**Fig 6.**
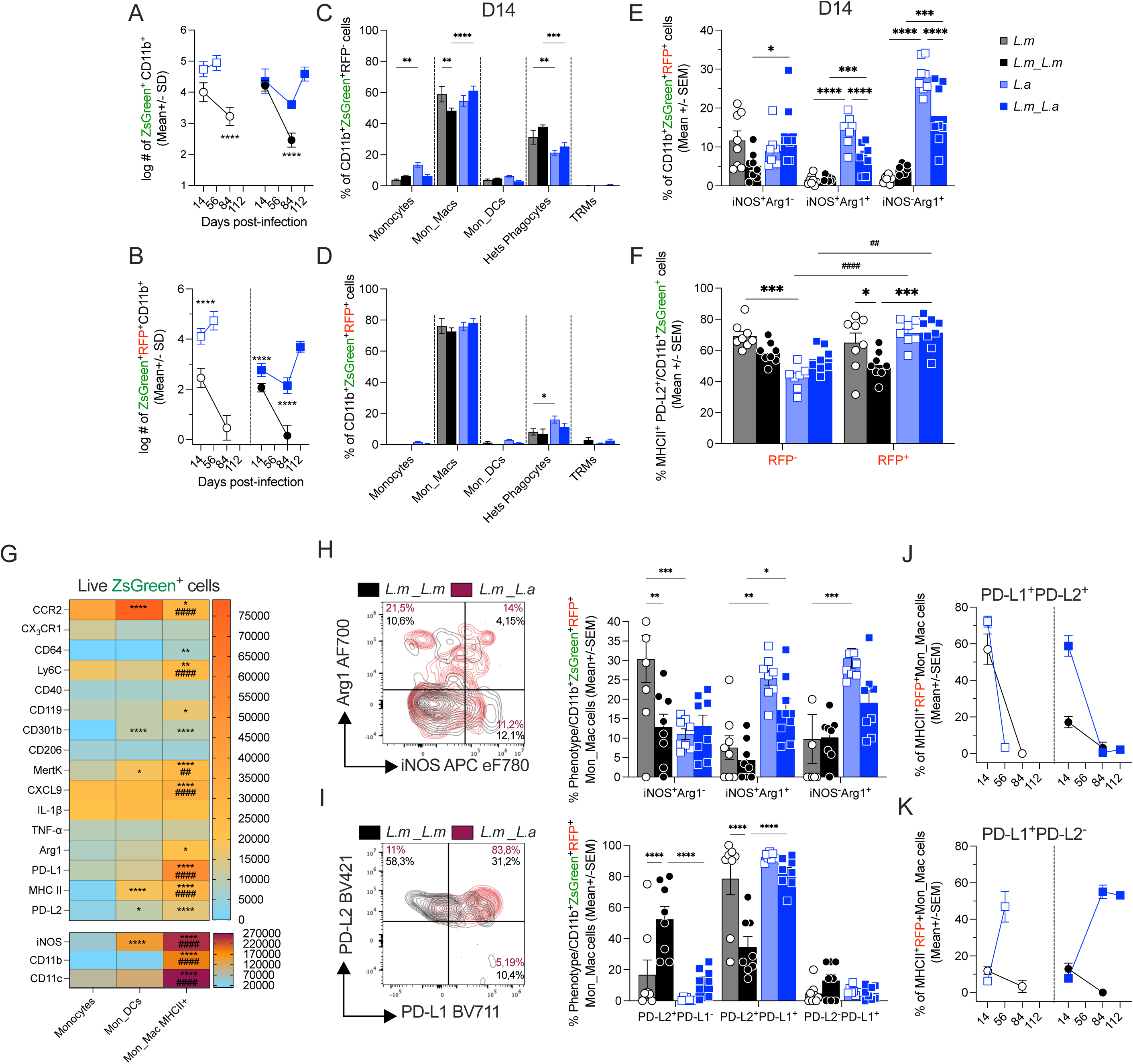
Monocyte fate tracing post-*Leishmania* challenge. Primary and secondary infections were performed as described in Figure 1 employing CCR2-creER^T2^.ZsGreen1 mice. Tmx treatment was performed after *Leishmania* challenge. The total #s of CD11b^+^ ZsGreen^+^ cells (A) or CD11b^+^ ZsGreen^+^ infected (RFP^+^) cells (B) and the % of the indicated cell subsets within RFP^-^CD11b^+^ZsGreen^+^ cells (C) or RFP^+^CD11b^+^ZsGreen^+^ cells (D) were determined by flow. The % of Arg1 and/or iNOS cells within the RFP^+^ZsGreen^+^CD11b^+^ population (E) and the % of MHCII^+^PD-L2^+^ within the RFP^+^ or RFP^-^ ZsGreen^+^CD11b^+^ population was determined by FLOW at D14 post-challenge (F). MFI heat-map for the indicated markers within the indicated ZsGreen^+^ subsets was determined at D14 post-challenge by FLOW (G). The % of Arg1 and/or iNOS cells (H) or PD-L1 and/or PD-L2 (I) within the RFP^+^Mon_Macs MHCII^+^ cells were determined by FLOW at D14 post-challenge. The kinetic dynamic of PD-L1^+^PD-L2^+^ (J) or PD-L1^+^PD-L2^-^ (K) RFP^+^ Mon_Macs MHCII+ cells at indicated time points were determined by FLOW. Data are from pooled experiments from 2 independent experiments. n=3-5 mice per time point per experiment. *p≤0.0381,**p≤0.0099,***p≤0.0003 ****≤0.0001.

Analysis of total ZsGreen^+^ cells at day 4 post-secondary challenge revealed large numbers of monocyte derived cells labelled prior to challenge at the site of infection, but over time (D14 and 42) their number and frequency declined, indicative of cell turnover at a site of infection induced inflammation (Fig 5, D and E). This initial wave of infiltrating CCR2^+^-monocytes differentiated primarily into CCR2^-^ Mon_Macs followed by CCR2^-^ Het_Phagocytes (Fig 5F, and Fig S3C). We also found large numbers (Fig 5G) and frequencies (Fig 5H) of RFP^+^ZsGreen^+^ infected cells. Remarkably, ZsGreen^+^ monocyte-derived cells accounted for approximately 80% of all infected RFP^+^CD11b^+^ cells in the skin on day 4 post-secondary challenge (Fig 5H). RFP^+^ infected monocyte-derived cells matured almost exclusively into Mon_Macs (Fig 5I and S3D), which no longer express CCR2 (S2A, **gating strategy**). Infected ZsGreen^+^ cells were significantly enriched for Mon_Macs versus uninfected cells at sites of secondary challenge, compare Fig. 5F with 5I (p≤0.001, n=6). However, no differences were observed when comparing *L.m._L.m.* versus *L.m._L.a.* infections. Intra-vital 2-photon microscopy confirmed ZsGreen^+^ cells infected with *Leishmania* parasites (Fig 5J) including cells with multiple phagosomal parasites, associated with enlarged phagolysosomes (yellow arrows), indicating replication, as well as parasite morphology, associated with reduced phagolysosomes size, a feature often associated with parasite killing (*67, 68*) (white arrows).

Having found no overt phenotypic differences in the phenotype of ZsGreen^+^ infected monocyte-derived cells recruited at the time of infection (Fig. 5I) we next determined if the highly infected Mon_Macs population possessed a permissive MHCII^+^ and PD-L2^+^ phenotype, as reported previously at primary sites of infection (Fig 5K)(*6*). Analysis of ZsGreen^+^ cells revealed increased frequencies of MHCII^+^PDL2^+^ cells within the RFP^+^ infected population on day 14 post-challenge, regardless of the parasite employed for challenge (Fig. 5K, right panel, RFP^+^ versus RFP^-^ populations, p≤0.0225, n≥11). In addition, secondary challenge sites with pre-existing Th1 III immunity had lower frequencies of MHCII^+^PD-L2^+^ infected cells versus primary sites, reflecting a more Th1-activated protective phenotype(*17*) and *L.m._L.a.* infections actually had a slightly lower frequency of MHCII^+^PD-L2^+^ Mono_Macs versus Mono_Macs from *L.m.*_*L.m.* challenge sites. Remarkably, despite the significant differences in the ultimate outcome of challenge infection (Fig 2; Fig 5, B and C), we also found no differences in Arg1 and iNOS production by infected RFP^+^CD11b^+^ZsGreen^+^ cells recruited at the time of secondary challenge in *L.m._L.m.* versus *L.m._L.a.* groups (Fig 5L). In addition, RFP^+^CD11b^+^ZsGreen^+^ cells expressed a predominant iNOS^+^Arg^-^ phenotype at D4 post-challenge, consistent with our analysis of total CD11b^+^ cells in Figure 3. Therefore, lineage tracing of the initial wave of recruited monocytes once again demonstrated early monocyte activation and maturation is intact in both challenge settings and is unlikely to be responsible for the loss of protective immunity. However, over time these cells no longer represented a large portion of the infected population (Fig. 5H), as expected given the cell turnover at an active site of infection. By D42 p.i., those RFP^+^ Mono_Macs that did remain were predominantly iNOS^-^Arg1^+^ (Fig. 5L), suggesting infected cells at latter time points may be exposed to a pro-parasitic environment regardless of overall parasite numbers (Fig 5C) and pre-existing Th1 immunity. In addition, we also found increased production of the monocyte recruiting chemokine CXCL9 at d42 p.i./ch. by infected monocyte derived cells (Fig 5M), reflecting our gene expression observations in Fig 2 and suggesting infected monocytes may drive the recruitment of new monocytic host cells into this environment in a positive feedback loop that may provide an ongoing source of phagocytic host cells(*6*) .

### CCR2-Lineage-Tracing reveals monocytes recruited at late timepoints mature into PD-L1^+^PD-L2^+^Arg^+^ macrophages and provide a unique immune-evasive parasitic niche

The late acquisition of an iNOS^-^Arg^+^ phenotype and chemokine production by infected monocyte-derived cells suggested monocytes recruited at later time points post-challenge may adopt a similar phenotype and provide the elusive hospitable and immune-evasive parasitic niche for *L.a.*. Therefore, we also Tmx treated mice as per Fig 5A, bottom row, to track ZsGreen^+^ monocyte-derived cells recruited to the site at different times *after* challenge. ZsGreen^+^ monocytes were recruited in significant numbers at all time points tested (Fig 6A), a large number were infected (Fig. 6B), and a significantly greater number were infected at *L.a.* versus *L.m.* challenge sites (Fig 6B). Both RFP^-^ uninfected (Fig 6C, and Fig S3E) and RFP^+^ infected (Fig 6D; Fig S3F) cells showed maturation towards a predominant Mon_Mac phenotype, which no longer express CCR2 (Fig 5K and Figs S2A and S3B) and this was more pronounced within the infected population (Fig 6, C versus D, p≤0.0013, n=8). Importantly, *L.a.* infected monocyte-derived cells recruited to sites of *L.a.* challenge on or just before day 14 post-challenge were drastically enriched for Arginase production compared to *L.m.*-infected cells (Fig 6E). In addition, infected RFP^+^ monocyte-derived cells at sites of *L.m._L.a.* challenge had higher MHCII^+^PD-L2^+^ frequencies versus those at sites of *L.m._L.m.* challenge (Fig. 6F, RFP+ closed blue squares versus closed black circles) and only *L.a* infected cells showed an enrichment of the MHCII^+^PD-L2^+^ phenotype versus RFP^-^ uninfected cells (Hash symbols in Fig 6F). The reduction in MHC II^+^PD-L2^+^ frequencies observed on ZsGreen^+^ infected monocytes in *L.m_L.m* versus *L.m.* mice was also lost in *L.m_L.a.* versus *L.a.* challenged mice (Fig 6F). This was in stark contrast to our day 14 analysis of cells that were recruited *prior* to challenge infection in Figure 5. Of interest, cells at sites of secondary *L.a.* challenge had lower frequencies of Arginase+ cells versus primary sites (Fig. 6E) once again demonstrating evidence of exposure to Th1 immunity. To further define these cells in the context of the other major phagocytic cells at sites of challenge we performed a heat-map median fluorescence intensity (MFI) analysis of ZsGreen^+^MHCII^+^ Mon_Macs comparing with monocytes and monocyte-derived DCs on day 14 post-challenge (Fig 6G). Our analysis revealed a highly defined and distinct population of cells that possess markers of both infection driven inflammation (CXCL9, iNOS, CD11c) and a tissue resolution/immuno-pathology response (PD-L1, PD-L2, Arg1, MertK), and the dual nature of these cells when infected with *L.a.* was emphasized by the significantly larger frequencies expressing both Arg and iNOS. Of note, these cells maintained very high levels of CD11b, the major receptor employed by phagocytes to phagocytose the *Leishmania* parasite(*69, 70*). In addition, most RFP^+^ Mon_Macs were Arg1^+^ (∼40%) (Fig 6H) and PD-L2^+^PD-L1^+^ (∼80%) (Fig 6I), contrasting with *L.m_L.m,* where only (∼15%) are Arg1^+^ and (∼30%) are PD-L2^+^PD-L1^+^. While most of infected Mon_macs are PD-L2^+^PD-L1^+^ cells at D14 (Fig 6J), it was PD-L1^+^ expression that was maintained at very late time points (Fig 6K). As PD-L1 is an immune-checkpoint inhibitor, that can be induced on dendritic cells infected with *L.a* and impair Th1 immunity and IFN-γ production(*71, 72*), this expression correlates with the surprisingly low activation levels of CD4 T cells at late time points in *L.a.* challenged mice despite high parasite loads (Fig. 2).

These observations implicate Arg1^+^MHCII^+^PD-L1^+^PD-L2^+^ monocyte-derived macrophages recruited at late time points as the elusive parasitic niche that drives the loss of protection following *L.a.* challenge.

### Anti PD-1 immune-checkpoint blockade enhances Th1 immunity but does not improve parasite killing after primary or secondary challenge

The high levels of PD-L1 and PD-L2 on *L.a.* infected monocyte-derived macrophages suggested PD-L1/2-PD-1 immune-checkpoint inhibition of Th1 immunity may explain the lack of protection following *L.a.* challenge. Therefore, we treated C57BL/6 mice with an anti-PD-1 antibody starting on d14 following primary or secondary challenge. Control mice received an isotype antibody, as described in the Methods. Treatments were administered twice weekly for four weeks, and mice were euthanized two days after the final dose, at d42 post-infection/challenge.

At day 42, approximately 60-75% of CD4^+^CD44^+^ T cells expressed PD-1 following primary infection, or ∼48% after secondary challenge (Fig 7A), confirming expression of this checkpoint inhibitor on T cells. Treatment did not alter the frequency PD-1^+^ cells (Fig. 7A) nor did it change the total # of CD4^+^CD44^+^ T cells in the skin (Fig 7B). However, treatment significantly enhanced the frequency of both CD119 (IFN-γ R1) expressing and IFN-γ producing CD4^+^ T cells in the skin upon *ex-vivo* antigen restimulation (Fig 7, C and D), indicating an augmented Th1 immune response. Consistent with this, anti-PD-1 blockade increased the frequency of iNOS (Fig. 7E) and CD119 (IFN-γ R1) (Fig. 7F) expressing cells among total mononuclear cells, without affecting Arg1 expression (Fig 7E). Specifically, this increase occurred in the iNOS^+^Arg1^-^ cells, with no change in the iNOS^+^Arg1^+^ population (Fig S4D and E). Despite this robust increase in Th1 immunity and phagocyte effector function, anti-PD-1 blockade resulted in reduced lesions size (Fig 7G and S4A) without affecting parasite burden (Fig 7H and S4B). This effect was consistent in both primary and secondary infections (Figs 7G, 7H, S4A and S4B). The decreased pathology was associated with reduced immune cell infiltration at the site of infection (Fig S4C). In an attempt to explain the disconnect between enhanced Th1 immunity and parasite load we also analyzed RFP^-^ uninfected versus RFP^+^ infected cells. Notably, the increase in iNOS (Fig. 7I) and CD119 (Fig. 7J) expression was restricted to only uninfected monocyte-derived macrophages and was not observed within infected cells. MHC II expression levels followed a similar pattern (Fig. 7K). These data suggest that *L.a* infected cells are refractory to anti-PD-1 immune-checkpoint blockade and that infection impairs the ability of host cells to further upregulate the IFN-γ receptor or MHC II in the presence of anti-PD-1-mediated enhanced Th1 immunity.

**Fig 7.**
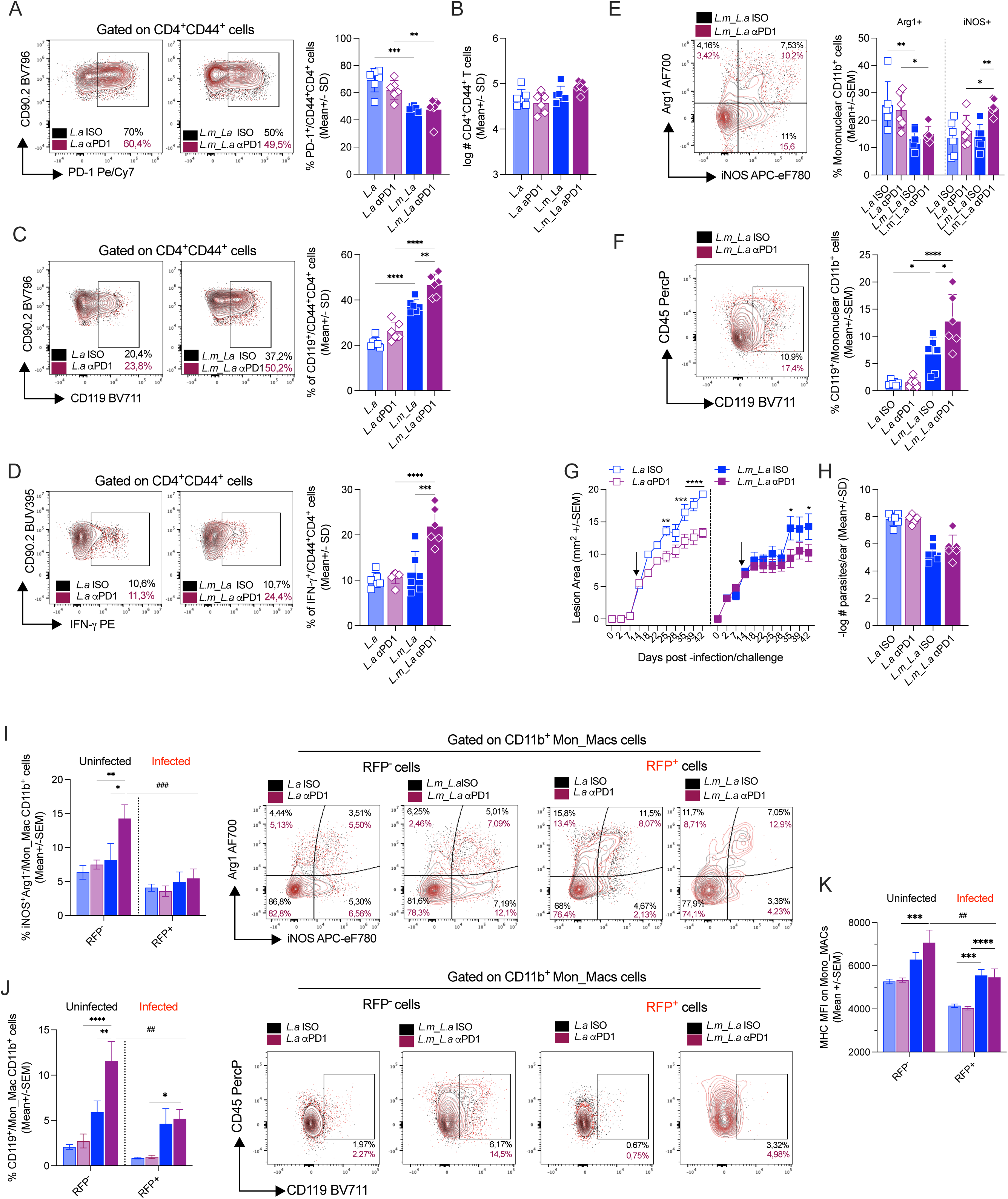
Infected Mon_Macs are refractory to enhanced Th1 immunity induced by PD1 blockade. Primary and secondary infections were performed as described in Figure 1. The %s of CD4^+^CD44^+^ cells expressing PD1 (A), CD119 (C) or IFN-γ (D) were determined by FLOW. The total #s of CD4^+^CD44^+^ were determined by FLOW 42 days post-infection challenge. The %s of mononuclear phagocytic cells expressing Arg1 and/or iNOS (E) or CD119 (F) were determined by FLOW 42 days post-infection challenge. Lesion area over-time (G) and #s of parasites (H) were determined at d42 post-primary and secondary infections. Mice start receiving anti-PD1 treatment or isotype control at the d14 post-infection/challenge with *L.a*.. The % within RFP- or RFP+ Mon-Macs expressing Arg1 or iNOS (I), CD119 (J), or MHC II (K) was determined by FLOW at d42 post-infection challenge. Data are from 1 representative of 2 independent experiments. n=3 mice per group. *p≤0.0214,**p≤0.0086,***p≤0.0003 ****≤0.0001.

Taken together, these data indicated that the loss of protection following *L.a* challenge is not directly mediated by PD-L1 or PD-L2 mediated suppression of Th1 immunity via PD-1 on T cells, as, even in the context of anti-PD-1, infected cells still fail to respond to the enhanced Th1 response. Rather, it reflects the parasites’ ability to modulate host cell responsiveness to Th1 immunity, specifically within infected monocyte-derived populations. These findings suggest that limiting the recruitment of inflammatory monocytes to the infection site may be a more effective strategy to control parasite growth by reducing the opportunity for parasite-mediated monocyte-derived host cell manipulation.

### Monocyte depletion following secondary *L.a.* challenge prevents progressive disease and reduces parasite numbers

To formally demonstrate the net pathogenic role for monocytes recruited at late time points in the loss of protective immunity provided by III, we employed CCR2-DTR mice, in which CCR2 expressing cells can be transiently depleted during the chronic stages of disease via timed administration of DT (*7, 73*). At chronic time points, infected monocyte-derived cells already present at the site of infection are Mon_Macs or heterogenous phagocytes and are CCR2-negative (Fig. 5 and 6). Therefore, DT treatment will target the circulating CCR2^+^ monocyte population, eliminating the recruitment of new monocytes to the site of infection and their availability to act as host cells for subsequent infection. Neutrophils, another inflammatory phagocytic cell infected by *Leishmania,* do not express CCR2 during cutaneous Leishmaniasis, so are not depleted (Fig. S5, A and B). Due to the vastly different infection kinetics in primary versus secondary infections, DT was administered at different times to attempt to reverse the non-healing phenotype at a relevant time point before infections reached their humane endpoint. Therefore, during primary *L.a.* infection, we started DT treatment after 6 wks and observed significant impact on lesion development after 5 DT treatments (Fig 8A), although parasite numbers were equivalent versus DTR^-^ littermate controls at the experimental endpoints, which were different (Fig 8B). During secondary challenge with *L.a* parasites, monocyte depletion resulted in smaller lesions (Fig 8C), a 50X reduction in parasite load by LDA (Fig 8D, note the log y-axis), reduced numbers of total infected RFP^+^CD11b^+^ cells (Fig 8E), fewer total (Fig S5C) and fewer infected monocyte derived cells (Fig 8F). To confirm our observations, we also employed the anti-CCR2 antibody MC-21(*47*) to deplete CCR2-monocytes over a 5-day period. Short-term monocyte depletion during the chronic stages of disease in both primary infection (Fig 8G and 8H), and secondary heterologous *L.a.* challenge, resulted in smaller lesions (Fig 8I), a 14X reduction in infected cell number by LDA (Fig. 8J) and fewer infected RFP^+^CD11b^+^ cells, 8K) per ear. As expected, total numbers of CCR2-expressing monocytes and monocyte-derived cells were reduced in the MC-21 treated group (Fig S5D), including infected monocyte-derived CCR2-negative Mon_DCs and Mon_Mac populations (Fig 8L). We also observed a reduction in the numbers of total dermal CD11b^+^CXCL9^+^ cells following both DT treatment (Fig 8M) and anti-CCR2 depletion (Fig 8N), suggesting that CCR2^+^ monocytes differentiate into the infected Arg^+^MHCII^+^PD-L1^+^PD-L2^+^CXCL9^+^ phenotype and that their removal reduces a monocyte-derived CXCL9 producing population. Because monocytes have important roles in tissue maintenance and healing, their long-term removal can cause tissue pathology and enhanced neutrophil recruitment, potentially confounding data interpretation(*7, 74, 75*), (*47*). Therefore, it is important to mention that we only depleted CCR2^+^ cells for a limited period and both of the CCR2-mediated monocyte depletion regimes we employed did not alter total or infected neutrophil numbers at the skin site after *Leishmania* challenge (Fig. S5B).

**Fig 8.**
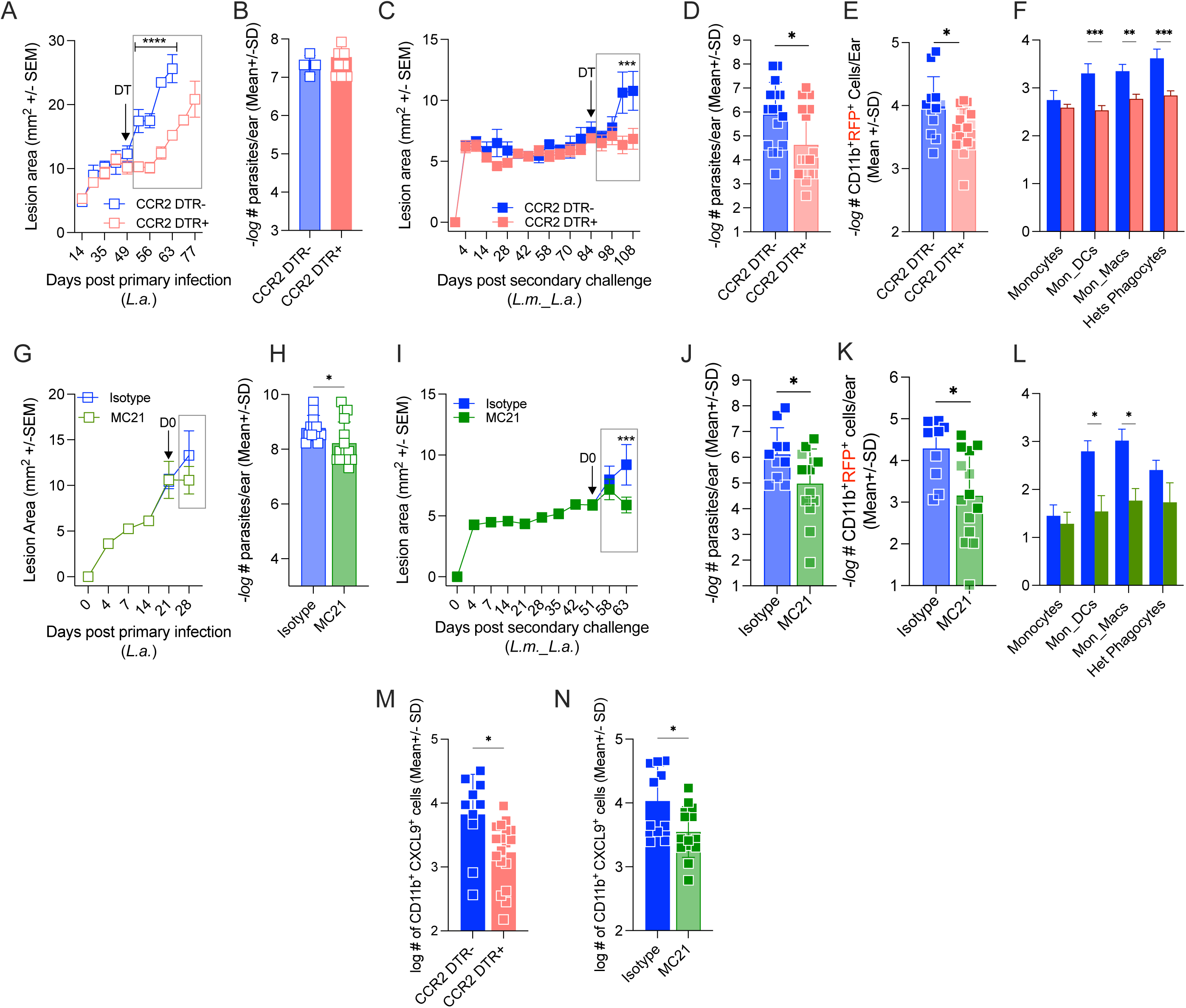
Monocyte depletion following secondary *L.a.* challenge prevents progressive disease and reduces parasite numbers. CCR2 DTR mice were treated with DT 5x every other day starting at the indicated DT timepoint. Mice were euthanized 2 days after the last DT dose. CCR2-DTR mice with a primary *L.a* infection received their first dose of DT at D49 and mice with a secondary *L.a* infection received their first DT dose at day 84 post-challenge. Lesion area over-time (A and C) and # of parasites (B and D) was determined at d77 post-primary infection or d108 post-secondary challenge by LDA. Total #s of infected CD11b^+^ cells (E) and the #s of indicated mononuclear subsets (F) within CD11b^+^RFP^+^ cells was determined by FLOW at d108 post-challenge. C57BL/6 mice were treated with CCR2 neutralizing antibodies MC-21 or its respective isotype control. Mice received the first dose of anti-CCR2 antibody (MC-21) at D21 (primary *L.a* infection) or at D51 (secondary *L.a* challenge). Mice were treated daily for five consecutive days. Lesion area over time (G and I) and # of parasites (H and J) were determined at d28 post-infection or d63 post-secondary challenge. Total #s of infected CD11b^+^ cells (K) and the #s of indicated mononuclear subsets (L) within CD11b^+^RFP^+^ cells were determined by FLOW at d63 post-challenge. (M) Total #s of CXCL9^+^CD11b^+^ cells following DT treatment as per (D-F) or (N) following anti-CCR2 as per (J-L). Data are from pooled experiments from ≥ 2 independent experiments. n=3-5 mice per time point per experiment. Darker and lighter shades of green/blue denote data from different independent experiments. *p≤0.019, **p≤0.0083, ***p≤0.0004, ****≤0.0001.

Taken together, our data suggest that CCR2-based depletion removes monocytes that are the precursors to CCR2-negative monocyte-derived phagocytes, thereby reducing their availability to act as host cells for subsequent infection and demonstrating their net pathogenic role following *L.a.* secondary challenge.

## Discussion

A major limitation to understanding how some intracellular pathogens employ the phagolysosome as a site of chronic infection and replication is defining the relevant host cells that provide this pathogen-niche. Infection with the *Leishmania* parasite represents a well-studied model of disease in this setting. Certain *Leishmania* strains are associated with more severe forms of cutaneous disease, such as *L. amazonensis* and *L. brasiliensis* in the new world, and *L.major*-Sd in the old word, which either require longer treatment or do not resolve(*76, 77*). Particularly, *L.a* and *L.m*-Sd parasites can thrive in the context of polarized Th1 immunity, challenging the well-established concept of a direct association between Th1 immunity and resolution (*78*). In the case of *L.a.*, these parasites appear to take advantage of the Th1-mediated inflammatory response including infection of recruited monocytic host cells (*79*) (*80, 81*) (*1, 50, 82*) (*6*). In contrast, we and others have previously shown that pre-existing Th1 III mediates rapid activation of monocytes following secondary challenge and results in robust protection from secondary *L. major* infections (*7, 59*). In the III setting, where monocytes are rapidly activated prior to the establishment of the parasitic niche, it would appear reasonable to assume the pre-existing response would provide protection against secondary challenge with more pathogenic strains. However, we showed that while pre-existing Th1 III was intact after heterologous challenge with *L. amazonensis*, it ultimately failed to provide protection.

Several lines of evidence suggest that this failure is not due to a deficiency in the elicitation of effective infection induced Th1 immunity: i) if protection were dependent upon unique *L.a.* strain-specific antigens, improved outcomes might be expected in the homologous challenge setting (*L.a_L.a*), but this was not the case, ii) while we did not define the T cell epitopes generated by the three *L.a.x L.m.* hybrids, the inheritance characteristics of *Leishmania* hybrids suggest proteins from each parental line would be expressed in the context of a healing phenotype at the primary infection site (Fig. 1G), yet III mediated by hybrids also failed to improve upon that provided by the parental *L.m.* parasite; iii) abrogation of potential immunoregulation of the Th1 response by Th2 immunity or IL-10 employing STAT6^-/-^ or IL-10^-/-^ deficient mice did not prevent the non-healing phenotype; iv) the number of *Leishmania*-specific CD4⁺ T cells as determined by quantification of PEPCK-tetramer positive cells or by analysis of cytokine-production by polyclonal T cells following *ex-vivo* antigen stimulation was indistinguishable at homologous versus heterologous challenge sites; v) enhancing Th1 immunity via anti-PD-1 immune checkpoint blockade did not result in reduced parasite loads, suggesting an infected-cell intrinsic immune-escape mechanism; vi) phagocyte activation, which ultimately is the effector mechanism for parasite control, was also equivalent when analyzed at day 4 post-challenge, a timepoint we have previously shown is critical for parasite control. Therefore, our data suggest the deficiency in *L.m.*-mediated III is not due to a defect in the pre-existing Th1 immune response.

Rather, employing careful time-dependent monocyte-lineage tracing, analysis of infected versus uninfected cells, and high-parameter spectral flow cytometry, we found that in stark contrast to those monocytes recruited at the time of challenge, monocytes recruited starting at approximately day 14 post *L.a.*-challenge acquired a Arg^+^MHCII^+^PD-L1^+^PD-L2^+^CXCL9^+^ phenotype that was distinct from the Arg^-^ dominant phenotype at sites of *L.m.*-FV1 or *L.m.-RYN* challenge, which were effectively controlled. Of note, *L.m*-Sd challenge was also not controlled, potentially due to preferential infection of tissue-resident macrophages that are also refractory to Th1 immunity(*9*) or other modes of evasion(*54, 83*)*. L.a.* infected Mono_Macs cells acquired a unique activation signature characterized by the expression of inflammatory (CXCL9, CD119 (IFN-γR1) and iNOS), phagocytic, (CD11b and CD11c), wound-healing (MertK and Arg1), and immune inhibitory checkpoint (PD-L1 and PD-L2) markers. PD-L1 is upregulated upon exposure to IFN-γ (*62*) and it’s expression on innate cells along with expression of the corresponding receptor PD-1 on T cells inhibits both T cell proliferation and cytokine production in peripheral tissues(*84, 85*). In the context of infectious diseases, the PD-L1/L2 – PD-1 immune checkpoint is associated with pathogen persistence during malaria, HIV and hepatitis B virus infections; and can impact Th1 priming during *Leishmania* infection(*71, 72, 85–87*). Therefore, we employed anti-PD-1 immune checkpoint blockade and predicted it would lead to increased Th1 immunity and reduced parasite loads. We found that this checkpoint was at least partially responsible for the low T cell responses observed following *L.a.* challenge despite high parasite loads as PD-1 blockade resulted in increased IFN-γ production and IFN-γR (CD119) expression by CD4+ T cells. This observation at least partially explains why the simultaneous lack of STAT-6 and IL-10 signaling has a minor impact on protection after *L.a* challenge, as this pathway is Th1, not Th2 or IL-10, driven.

Immune-checkpoint blockade (anti-PD1) also enhanced iNOS and CD119, but not arginase, expression within the mononuclear phagocytic cell compartment, leading to decreased pathology, which was associated with a small but significant decrease in CD11b^+^ cells in the tissue. However, this was unliked to a reduction in parasite burden. Again, employing high-parameter spectral flow cytometry and tracking of infected cells, we were able to reveal that this enhancement was restricted to uninfected cells. *L.a.* infected CCR2-negative Mon_Macs, which account for the vast majority of infected cells, were refractory to PD-1 immune checkpoint blockade, failing to show enhanced iNOS or IFN-γR (CD119) expression and maintenance of parasite numbers. The ability of *L.a.* to survive in these cells, even in a setting of enhanced Th1 immunity may be the consequence of the selection of highly resistant parasites by the early pre-existing Th1 immune response(*88*). The collective findings indicate that these cells represent a specialized infection-induced macrophage state that remains refractory to IFN-γ-mediated activation despite the blockade of several suppressive pathways, including STAT6/Arg1 signaling, IL-10, and PD-1/PD-L1. Therefore, while PD-L1/L2 and Arginase expression is a property of these cells, it is not the causative factor for their refractory phenotype.

Lastly, to demonstrate that the net impact of late monocyte recruitment enhances diseases we depleted the CCR2^+^ cells that undergo maturation to become CCR2^-^ infected Mon_Macs. At this time-point, infected cells in the tissue are CCR2^-^ and therefore our CCR2-dependent depletion methodologies target the precursors to our cells of interest. As predicted based on a pathogenic role for late monocyte recruitment, transient monocyte-depletion resulted in a rapid reduction in total monocyte-derived cell numbers, lesion size, parasite loads and numbers of infected cells. This result is surprising given the protective role previously ascribed to monocytes in the context of pre-existing Th1 immunity at secondary sites of challenge(*7, 59*). These cells are distinct from monocyte-derived cells at primary chronic sites of infection in that they express significantly lower frequencies of Arginase^+^ cells but still fail to control parasite growth, supporting a limited role for the STAT6/Th2/Arginase axis as shown in Figure 4. Our observations support a model whereby late monocyte recruitment drives parasite expansion via both provision of permissive host cells and cell intrinsic regulation of subsequently infected Mono_Macs. Infection was not an absolute requirement to generate the Arg^+^MHCII^+^PD-L1^+^PD-L2^+^CXCL9^+^ phenotype as these cells also existed in the uninfected population. Having now identified these cells of interest, future studies can begin to address the degree to which *L.a.* targets these cells for infection or facilitates their generation and/or persistence.

Our results address the roles of Th1 immunity, immunoregulation, parasite species, host-cell phenotype, and most importantly the timing of monocyte recruitment in the pathogenesis of resistance of a phagosomal parasite to pre-existing Th1 immunity. Our finding do differ somewhat from those of Vanloubbeeck and Jones(*89*), but we would argue that the 5×10^6^ dose of non-purified stationary phase promastigotes at a s.c. challenge site in the footpad employed in this study may not sufficiently reflect physiological challenge infection.

Our results have important implications for both therapeutic and vaccine strategies which often operate under the assumption that induction of the optimal Th1 immune response will be sufficient to provide protection at a site of secondary challenge(*50*). Our findings support a model in which disease outcome is determined less by the magnitude of Th1 immunity and more by the availability and responsiveness of an intracellular pathogen-niche that remains refractory to effector activation. This uncoupling of Th1 immunity from host-cell antimicrobial function suggests successful treatment strategies against *L.a.* are more likely to be those that simultaneously target both the induction of robust Th1 immunity but also the intracellular biology of the parasite-phagocyte interaction. The work presented here defines the functional characteristics and specific host cell that needs to be targeted by these strategies and provides a direction for future investigation into the unique cellular biology of *L.a.* infection of monocyte-derived cells.

## Acknowledgments

This work was supported by Canadian Institutes of Health Research grant CIHR Project Grant 519427 to N.C.P., S.A.E.S. obtained a Brazilian CAPES fellowship (PDSE 2023). L.S.H. obtained graduate student salary support from the Department of Comparative Biology and Experimental Medicine, University of Calgary School of Veterinary Medicine. C.G received a Mitacs Globalink Graduate Fellowship Award. M.B.C. and L.S.H. received professional development training and travel funding from the University of Calgary Host-Parasite Interactions Training Program. We thank Dr. Karen Poon at the Nicole Perkins Microbial Communities Core Labs. We thank the NIH Tetramer Core Facility for providing the tetramer used in this work.

---

**Fig S1. Generating Infection Induced Immunity**. C57BL/6 mice were infected with 10^4^ self-healing *L. major*-Fv1 parasites in the footpad and lesion thickness was assessed over 20 wks post-infection (A). The red arrow indicated acquisition of the healed phenotype. Parasite load quantification in the footpad 20 wks post-infection with self-healing *L.m.-* Fv1 (B). (C-D) Concomitant immunity in Th2-prone, susceptible, BALB/c mice mediates a level of protection at a secondary i.d. challenge site in the ear skin despite progressive disease at the primary *L.m.* infection site in the footpad. BALB/c mice were infected with 10^4^self-healing *L.m.*-Fv1 parasites in the footpad and lesion thickness was assessed over 6 wks post-infection (C). Mice were then challenged with 2×10^5^ *L.m.* in the ear dermis and lesion size and parasite load quantification was performed (D). Data are pooled 4 independent experiments (A), or from 1 individual experiment (B-D), representative of 2 repeat experiments. n=3-5 mice per time point per experiment. ***p≤0.0003

**Fig S2. Gating strategy and innate cell infiltration.** Gating strategy employed to define the innate CD11b^+^ populations and T cells used in the study (A). Representative CD40, PD-L1, PD-L2 and MHC II do plots and overlay on the indicated groups at D4 p.i. (B). The total #s (C) of mononuclear phagocytic indicated cell subsets was determined by FLOW. Data are from pooled experiments from ≤ 3 independent experiments (C), or from individual experiments (A-B) representative of ≥ 2 repeat experiments. n=3-5 mice per time point per experiment. *p≤0.013,**p≤0.0082,***p≤0.0003 ****≤0.0001.

**Fig S3. Monocyte fate tracing pre or post *Leishmania* challenge.** Total #s of CD11b+ cells (A) and the indicated mononuclear phagocytic cells (B) at indicated time points were determined by FLOW. Cells were labelled either prior (C and D) or after (E and F) the infection/challenge with *Leishmania* parasites. The % of indicated cell subsets within total CD11b^+^ZsGreen^+^ cells (C and E) or within infected CD11b^+^ZsGreen^+^ cells (D and F) were determined by FLOW at indicated time points after infection/challenge. Data are pooled from 2 independent experiments. n=3-5 mice per time point per experiment.***p≤0.001 ****≤0.0001.

**Fig S4. Anti PD-1 immune-checkpoint blockade does not improve parasite killing after primary or secondary challenge.** C57BL/6 mice were treated with anti-PD1 or isotype control as described in Figure 7. Lesion area over-time (A) and # of parasites (B) were determined at d42 post-infection/challenge. The total #s of CD11b+ cells and the indicated cell subsets were determined by FLOW 42 days post-infection/challenge. The %s of iNOS/Ag1 within RFP^-^ (D) or RFP^+^ (E) Mon_Macs or the MHC II MFI on the Mono_DC population were determined by FLOW. CD119 expression on the indicated populations as described for Fig. 7J. Data are from 1 representative of 2 independent experiments. n=3 mice per group. *p≤0.0452,**p≤0.0011,***p≤0.0002 ****≤0.0001.

**Fig S5. Monocyte depletion following secondary *L.a.* challenge prevents progressive disease and reduces parasite numbers.** CCR2 DTR mice were treated as described in Figure 8. Mice were euthanized 2 days after the last DT dose. Flow plots indicating that Tmx treatment of CCR2-creER^T2^.ZsGreen1 mice does not labelled neutrophils after *Leishmania* infection (A) and depletion of CCR2+ cells does not impact neutrophils numbers in the skin (B) were determined by FLOW. The #s of indicated cell subsets (C) were determined by FLOW at d108 post-challenge. C57BL/6 mice were treated with CCR2 neutralizing antibodies MC-21 or its respective isotype control. Mice with a secondary *L.a* infection received the first dose of MC-21 at D63 and were treated daily for five consecutive days The #s of indicated cell subsets (D) were determined by FLOW at D77 post-challenge. Data are from individual experiments representative of ≥ 2 repeat experiments. n=3-5 mice per time point per experiment. *p≤0.016, ***p≤0.0004 ****≤0.0001.

## References

1. N. C. Peters, N. Khan, C. H. Mody, Novel approaches to preventing phagosomal infections: timing is key. Trends Immunol 44, 22–31 (2023).

2. M. Guilliams, A. Mildner, S. Yona, Developmental and Functional Heterogeneity of Monocytes. Immunity **49**, 595–613 (2018).

3. C. Shi, E. G. Pamer, Monocyte recruitment during infection and inflammation. Nat Rev Immunol 11, 762–774 (2011).

4. M. Guilliams, G. R. Thierry, J. Bonnardel, M. Bajenoff, Establishment and Maintenance of the Macrophage Niche. Immunity 52, 434–451 (2020).

5. N. C. Peters et al., In vivo imaging reveals an essential role for neutrophils in leishmaniasis transmitted by sand flies. Science 321, 970–974 (2008).

6. M. B. Carneiro et al., Th1-Th2 Cross-Regulation Controls Early Leishmania Infection in the Skin by Modulating the Size of the Permissive Monocytic Host Cell Reservoir. Cell Host Microbe, (2020).

7. A. Romano et al., Divergent roles for Ly6C+CCR2+CX3CR1+ inflammatory monocytes during primary or secondary infection of the skin with the intra-phagosomal pathogen Leishmania major. PLoS Pathog 13, e1006479 (2017).

8. S. Heyde et al., CD11c-expressing Ly6C+CCR2+ monocytes constitute a reservoir for efficient Leishmania proliferation and cell-to-cell transmission. PLoS Pathog 14, e1007374 (2018).

9. S. H. Lee et al., Mannose receptor high, M2 dermal macrophages mediate nonhealing Leishmania major infection in a Th1 immune environment. J Exp Med 215, 357–375 (2018).

10. G. Pessenda et al., Kupffer cell and recruited macrophage heterogeneity orchestrate granuloma maturation and hepatic immunity in visceral leishmaniasis. Nat Commun 16, 3125 (2025).

11. P. Scott, P. Natovitz, R. L. Coffman, E. Pearce, A. Sher, Immunoregulation of cutaneous leishmaniasis. T cell lines that transfer protective immunity or exacerbation belong to different T helper subsets and respond to distinct parasite antigens. J Exp Med 168, 1675–1684 (1988).

12. S. J. Green, M. S. Meltzer, J. B. Hibbs, Jr., C. A. Nacy, Activated macrophages destroy intracellular Leishmania major amastigotes by an L-arginine-dependent killing mechanism. J Immunol 144, 278–283 (1990).

13. R. Olekhnovitch, B. Ryffel, A. J. Muller, P. Bousso, Collective nitric oxide production provides tissue-wide immunity during Leishmania infection. J Clin Invest 124, 1711–1722 (2014).

14. C. De Trez et al., iNOS-producing inflammatory dendritic cells constitute the major infected cell type during the chronic Leishmania major infection phase of C57BL/6 resistant mice. PLoS Pathog 5, e1000494 (2009).

15. M. M. Kane, D. M. Mosser, The role of IL-10 in promoting disease progression in leishmaniasis. J Immunol 166, 1141–1147 (2001).

16. A. L. Dent, T. M. Doherty, W. E. Paul, A. Sher, L. M. Staudt, BCL-6-deficient mice reveal an IL-4-independent, STAT6-dependent pathway that controls susceptibility to infection by Leishmania major. J Immunol 163, 2098–2103 (1999).

17. M. B. Carneiro et al., Th1-Th2 Cross-Regulation Controls Early Leishmania Infection in the Skin by Modulating the Size of the Permissive Monocytic Host Cell Reservoir. Cell host & microbe 27, 752–768 e757 (2020).

18. R. B. Kennedy, I. G. Ovsyannikova, R. M. Jacobson, G. A. Poland, The immunology of smallpox vaccines. Curr Opin Immunol 21, 314–320 (2009).

19. C.-F. Team, Past SARS-CoV-2 infection protection against re-infection: a systematic review and meta-analysis. Lancet 401, 833–842 (2023).

20. S. K. Hoiseth, B. A. Stocker, Aromatic-dependent Salmonella typhimurium are non-virulent and effective as live vaccines. Nature 291, 238–239 (1981).

21. P. Mastroeni, B. Villarreal-Ramos, C. E. Hormaeche, Adoptive transfer of immunity to oral challenge with virulent salmonellae in innately susceptible BALB/c mice requires both immune serum and T cells. Infect Immun 61, 3981–3984 (1993).

22. L. R. Joslyn, J. L. Flynn, D. E. Kirschner, J. J. Linderman, Concomitant immunity to M. tuberculosis infection. Sci Rep 12, 20731 (2022).

23. F. L. Wormley, Jr., J. R. Perfect, C. Steele, G. M. Cox, Protection against cryptococcosis by using a murine gamma interferon-producing Cryptococcus neoformans strain. Infect Immun 75, 1453–1462 (2007).

24. M. S. Amaral et al., Rhesus macaques self-curing from a schistosome infection can display complete immunity to challenge. Nat Commun 12, 6181 (2021).

25. K. Obata-Ninomiya et al., The skin is an important bulwark of acquired immunity against intestinal helminths. J Exp Med 210, 2583–2595 (2013).

26. D. L. Doolan, C. Dobano, J. K. Baird, Acquired immunity to malaria. Clin Microbiol Rev 22, 13–36, Table of Contents (2009).

27. N. C. Peters et al., Vector transmission of leishmania abrogates vaccine-induced protective immunity. PLoS Pathog 5, e1000484 (2009).

28. D. L. Sacks, Vaccines against tropical parasitic diseases: a persisting answer to a persisting problem. Nat Immunol 15, 403–405 (2014).

29. V. Sergiev et al., Epidemiology and Control of Leishmaniasis in the Former USSR: A Review Article. Iran J Parasitol 13, 342–350 (2018).

30. N. C. Peters et al., Chronic parasitic infection maintains high frequencies of short-lived Ly6C+CD4+ effector T cells that are required for protection against re-infection. PLoS Pathog 10, e1004538 (2014).

31. L. S. Hohman et al., Protective CD4+ Th1 cell-mediated immunity is reliant upon execution of effector function prior to the establishment of the pathogen niche. PLoS Pathog 17, e1009944 (2021).

32. N. D. Glennie, S. W. Volk, P. Scott, Skin-resident CD4+ T cells protect against Leishmania major by recruiting and activating inflammatory monocytes. Plos Pathogens 13, (2017).

33. K. S. Tabbara et al., Conditions influencing the efficacy of vaccination with live organisms against Leishmania major infection. Infection and immunity 73, 4714–4722 (2005).

34. N. C. Peters et al., Evaluation of recombinant Leishmania polyprotein plus glucopyranosyl lipid A stable emulsion vaccines against sand fly-transmitted Leishmania major in C57BL/6 mice. J Immunol **189**, 4832–4841 (2012).

35. H. Mahmoudzadeh-Niknam, G. Khalili, F. Abrishami, A. Najafy, V. Khaze, The Route of Leishmania tropica Infection Determines Disease Outcome and Protection against Leishmania major in BALB/c Mice. Korean Journal of Parasitology 51, 69–74 (2013).

36. A. Barral et al., Leishmaniasis in Bahia, Brazil: evidence that Leishmania amazonensis produces a wide spectrum of clinical disease. Am J Trop Med Hyg 44, 536–546 (1991).

37. M. B. Carneiro et al., IFN-gamma-Dependent Recruitment of CD4(+) T Cells and Macrophages Contributes to Pathogenesis During Leishmania amazonensis Infection. J Interferon Cytokine Res 35, 935–947 (2015).

38. N. Wanasen, C. L. MacLeod, L. G. Ellies, L. Soong, L-arginine and cationic amino acid transporter 2B regulate growth and survival of Leishmania amazonensis amastigotes in macrophages. Infect Immun 75, 2802–2810 (2007).

39. M. M. Chaves et al., The role of dermis resident macrophages and their interaction with neutrophils in the early establishment of Leishmania major infection transmitted by sand fly bite. PLoS Pathog 16, e1008674 (2020).

40. F. A. Neva, D. Wyler, T. Nash, Cutaneous leishmaniasis--a case with persistent organisms after treatment in presence of normal immune response. Am J Trop Med Hyg 28, 467–471 (1979).

41. M. B. H. Carneiro et al., NOX2-Derived Reactive Oxygen Species Control Inflammation during Leishmania amazonensis Infection by Mediating Infection-Induced Neutrophil Apoptosis. J Immunol 200, 196–208 (2018).

42. G. F. Spath, S. M. Beverley, A lipophosphoglycan-independent method for isolation of infective Leishmania metacyclic promastigotes by density gradient centrifugation. Exp Parasitol **GG**, 97–103 (2001).

43. E. Inbar et al., Whole genome sequencing of experimental hybrids supports meiosis-like sexual recombination in Leishmania. PLoS Genet 15, e1008042 (2019).

44. A. Romano et al., Cross-species genetic exchange between visceral and cutaneous strains of Leishmania in the sand fly vector. Proc Natl Acad Sci U S A 111, 16808–16813 (2014).

45. A. J. Pagan et al., Tracking antigen-specific CD4+ T cells throughout the course of chronic Leishmania major infection in resistant mice. Eur J Immunol 43, 427–438 (2013).

46. M. B. Carneiro, L. S. Hohman, J. G. Egen, N. C. Peters, Use of two-photon microscopy to study Leishmania major infection of the skin. Methods, (2017).

47. M. Mack et al., Expression and characterization of the chemokine receptors CCR2 and CCR5 in mice. J Immunol 166, 4697–4704 (2001).

48. A. Romano, N. A. Doria, J. Mendez, D. L. Sacks, N. C. Peters, Cutaneous Infection with Leishmania major Mediates Heterologous Protection against Visceral Infection with Leishmania infantum. J Immunol, (2015).

49. N. Kimblin et al., Quantification of the infectious dose of Leishmania major transmitted to the skin by single sand flies. Proc Natl Acad Sci U S A 105, 10125–10130 (2008).

50. L. S. Hohman, N. C. Peters, CD4(+) T Cell-Mediated Immunity against the Phagosomal Pathogen Leishmania: Implications for Vaccination. Trends Parasitol 35, 423–435 (2019).

51. C. Z. Gonzalez-Lombana et al., Early infection with Leishmania major restrains pathogenic response to Leishmania amazonensis and parasite growth. Acta Trop 106, 27–38 (2008).

52. P. Veras et al., A dhfr-ts-Leishmania major knockout mutant cross-protects against Leishmania amazonensis. Mem Inst Oswaldo Cruz **94**, 491–496 (1999).

53. E. Gorelik, Concomitant tumor immunity and the resistance to a second tumor challenge. Adv Cancer Res **39**, 71–120 (1983).

54. C. F. Anderson, S. Mendez, D. L. Sacks, Nonhealing infection despite Th1 polarization produced by a strain of Leishmania major in C57BL/6 mice. J Immunol 174, 2934–2941 (2005).

55. C. M. C. Catta-Preta, D. L. Sacks, Genetic Exchange in Leishmania: Understanding the Cryptic Sexual Cycle. Annu Rev Microbiol **79**, 105–128 (2025).

56. L. W. Stamper et al., Infection parameters in the sand fly vector that predict transmission of Leishmania major. PLoS Negl Trop Dis 5, e1288 (2011).

57. S. H. Lee, et al., M2-like, dermal macrophages are maintained via IL-4/CCL24-mediated cooperative interaction with eosinophils in cutaneous leishmaniasis. *Sci Immunol* 5, (2020).

58. A. Romano, N. A. Doria, J. Mendez, D. L. Sacks, N. C. Peters, Cutaneous Infection with Leishmania major Mediates Heterologous Protection against Visceral Infection with Leishmania infantum. J Immunol 195, 3816–3827 (2015).

59. N. D. Glennie, S. W. Volk, P. Scott, Skin-resident CD4+ T cells protect against Leishmania major by recruiting and activating inflammatory monocytes. PLoS Pathog 13, e1006349 (2017).

60. Z. Mou et al., Identification of broadly conserved cross-species protective Leishmania antigen and its responding CD4+ T cells. Sci Transl Med 7, 310ra167 (2015).

61. S. Tamoutounour et al., Origins and functional specialization of macrophages and of conventional and monocyte-derived dendritic cells in mouse skin. Immunity **3G**, 925–938 (2013).

62. P. Loke, J. P. Allison, PD-L1 and PD-L2 are differentially regulated by Th1 and Th2 cells. Proc Natl Acad Sci U S A 100, 5336–5341 (2003).

63. V. Iniesta et al., Arginase I induction during Leishmania major infection mediates the development of disease. Infect Immun 73, 6085–6090 (2005).

64. U. Schleicher et al., TNF-Mediated Restriction of Arginase 1 Expression in Myeloid Cells Triggers Type 2 NO Synthase Activity at the Site of Infection. Cell Rep 15, 1062–1075 (2016).

65. Y. Belkaid et al., The role of interleukin (IL)-10 in the persistence of Leishmania major in the skin after healing and the therapeutic potential of anti-IL-10 receptor antibody for sterile cure. J Exp Med 194, 1497–1506 (2001).

66. R. Zayats et al., Antigen recognition reinforces regulatory T cell mediated Leishmania major persistence. Nat Commun 14, 8449 (2023).

67. M. Rabinovitch et al., Destruction of Leishmania mexicana amazonensis amastigotes within macrophages in culture by phenazine methosulfate and other electron carriers. J Exp Med 155, 415–431 (1982).

68. C. C. Pessoa et al., ATP6V0d2 controls Leishmania parasitophorous vacuole biogenesis via cholesterol homeostasis. PLoS Pathog 15, e1007834 (2019).

69. A. J. Ranson et al., C3/CD11b-Mediated Leishmania major Internalization by Neutrophils Induces Intraphagosomal NOX2-Mediated Respiratory Burst but Fails to Eliminate Parasites and Induces a State of Stalled Apoptosis. J Immunol 211, 103–117 (2023).

70. D. M. Mosser, P. J. Edelson, The mouse macrophage receptor for C3bi (CR3) is a major mechanism in the phagocytosis of Leishmania promastigotes. J Immunol 135, 2785–2789 (1985).

71. A. M. da Fonseca-Martins et al., Immunotherapy using anti-PD-1 and anti-PD-L1 in Leishmania amazonensis-infected BALB/c mice reduce parasite load. Sci Rep **9**, 20275 (2019).

72. H. L. de Matos Guedes et al., Leishmania amazonensis infection induces PD-L1 expression on dendritic cells and impairs Th1 responses in vitro and in vivo. Sci Rep 15, 37856 (2025).

73. T. M. Hohl et al., Inflammatory monocytes facilitate adaptive CD4 T cell responses during respiratory fungal infection. Cell Host Microbe 6, 470–481 (2009).

74. J. R. Grainger et al., Inflammatory monocytes regulate pathologic responses to commensals during acute gastrointestinal infection. Nat Med **19**, 713–721 (2013).

75. R. M. Kratofil et al., A monocyte-leptin-angiogenesis pathway critical for repair post-infection. Nature **609**, 166–173 (2022).

76. F. T. Silveira, R. Lainson, C. E. Corbett, Clinical and immunopathological spectrum of American cutaneous leishmaniasis with special reference to the disease in Amazonian Brazil: a review. Mem Inst Oswaldo Cruz **GG**, 239–251 (2004).

77. F. T. Silveira, R. Lainson, C. M. De Castro Gomes, M. D. Laurenti, C. E. Corbett, Immunopathogenic competences of Leishmania (V.) braziliensis and L. (L.) amazonensis in American cutaneous leishmaniasis. Parasite Immunol 31, 423–431 (2009).

78. P. Scott, F. O. Novais, Cutaneous leishmaniasis: immune responses in protection and pathogenesis. Nat Rev Immunol 16, 581–592 (2016).

79. M. B. Carneiro, N. C. Peters, The Paradox of a Phagosomal Lifestyle: How Innate Host Cell-Leishmania amazonensis Interactions Lead to a Progressive Chronic Disease. Front Immunol 12, 728848 (2021).

80. L. M. Sousa et al., IL-18 contributes to susceptibility to Leishmania amazonensis infection by macrophage-independent mechanisms. Cytokine 74, 327–330 (2015).

81. L. Soong et al., Role of CD4+ T cells in pathogenesis associated with Leishmania amazonensis infection. J Immunol 158, 5374–5383 (1997).

82. S. H. Lee et al., Dermis resident macrophages orchestrate localized ILC2 eosinophil circuitries to promote non-healing cutaneous leishmaniasis. Nat Commun 14, 7852 (2023).

83. M. Charmoy et al., The Nlrp3 inflammasome, IL-1beta, and neutrophil recruitment are required for susceptibility to a nonhealing strain of Leishmania major in C57BL/6 mice. Eur J Immunol 46, 897–911 (2016).

84. M. E. Keir, M. J. Butte, G. J. Freeman, A. H. Sharpe, PD-1 and its ligands in tolerance and immunity. Annu Rev Immunol 26, 677–704 (2008).

85. M. N. Wykes, S. R. Lewin, Immune checkpoint blockade in infectious diseases. Nat Rev Immunol 18, 91–104 (2018).

86. N. S. Butler et al., Therapeutic blockade of PD-L1 and LAG-3 rapidly clears established blood-stage Plasmodium infection. Nat Immunol 13, 188–195 (2011).

87. R. A. Zander et al., PD-1 Co-inhibitory and OX40 Co-stimulatory Crosstalk Regulates Helper T Cell Differentiation and Anti-Plasmodium Humoral Immunity. Cell Host Microbe 17, 628–641 (2015).

88. M. S. da Silva et al., Consequences of acute oxidative stress in Leishmania amazonensis: From telomere shortening to the selection of the fittest parasites. Biochim Biophys Acta Mol Cell Res 1864, 138–150 (2017).

89. Y. Vanloubbeeck, D. E. Jones, Protection of C3HeB/FeJ mice against Leishmania amazonensis challenge after previous Leishmania major infection. Am J Trop Med Hyg 71, 407–411 (2004).

