## Supplemental Figures for "Pre-existing Th1 immunity is abrogated by ongoing recruitment of monocytic host cells that are refractory to activation"

**Fig S5. Monocyte depletion following secondary *L.a.* challenge prevents progressive disease and reduces parasite numbers.** CCR2 DTR mice were treated as described in Figure 8. Mice were euthanized 2 days after the last DT dose. Flow plots indicating that Tmx treatment of CCR2-creER$^{T2}$.ZsGreen1 mice does not labelled neutrophils after *Leishmania* infection (A) and depletion of CCR2+ cells does not impact neutrophils numbers in the skin (B) were determined by FLOW. The #s of indicated cell subsets (C) were determined by FLOW at d108 post-challenge. C57BL/6 mice were treated with CCR2 neutralizing antibodies MC-21 or its respective isotype control. Mice with a secondary *L.a* infection received the first dose of MC-21 at D63 and were treated daily for five consecutive days The #s of indicated cell subsets (D) were determined by FLOW at D77 post-challenge. Data are from individual experiments representative of ≥ 2 repeat experiments. n=3-5 mice per time point per experiment. *p≤0.016, ***p≤0.0004 ****≤0.0001.

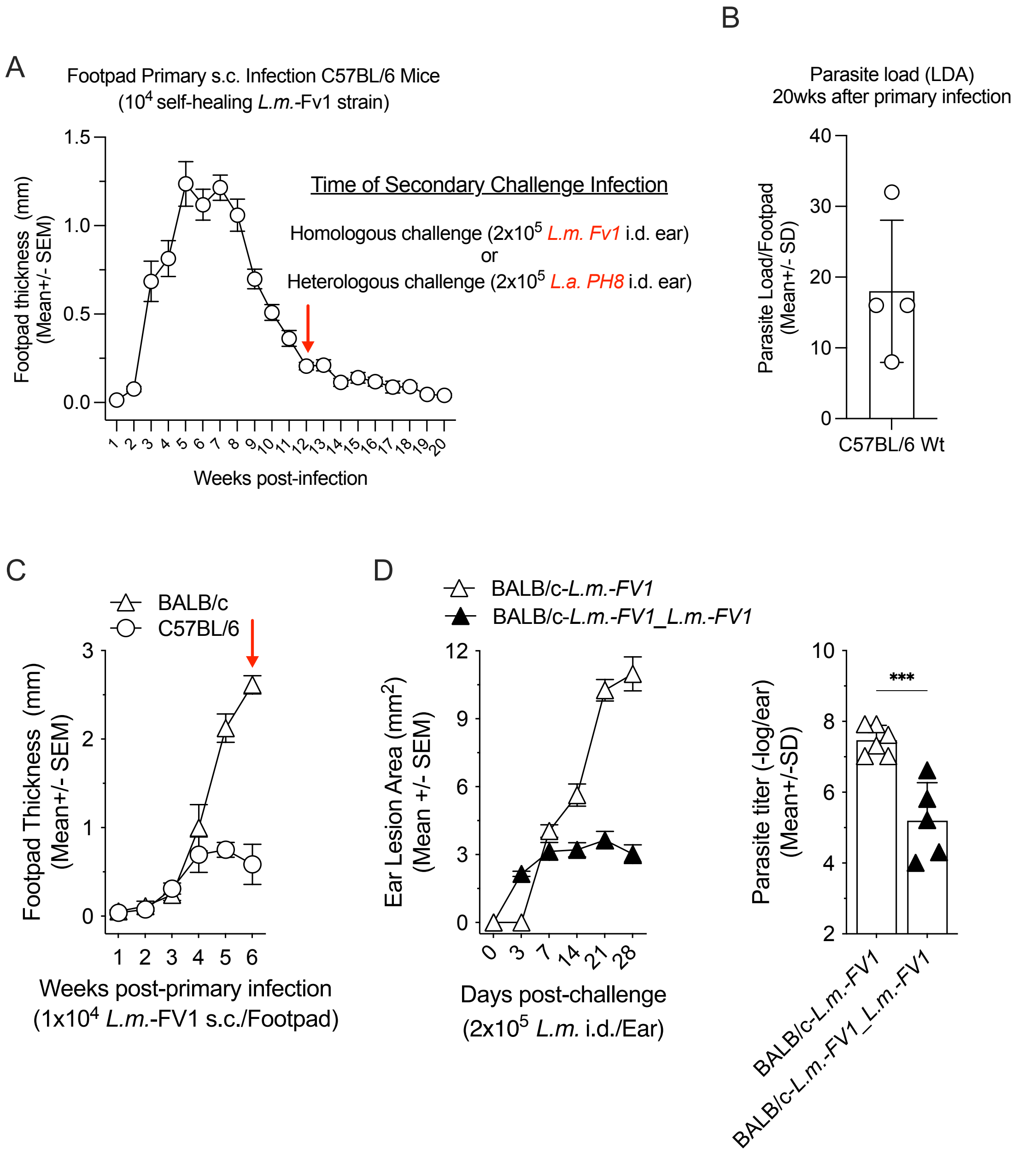
A
Footpad Primary s.c. Infection C57BL/6 Mice
(10^4 self-healing L.m.-Fv1 strain)
Footpad thickness (mm) (Mean+/- SEM)
Weeks post-infection
Time of Secondary Challenge Infection
Homologous challenge (2x10^5 L.m. Fv1 i.d. ear)
or
Heterologous challenge (2x10^5 L.a. PH8 i.d. ear)
B
Parasite load (LDA)
20wks after primary infection
Parasite Load/Footpad (Mean+/- SD)
C57BL/6 Wt
C
BALB/c
C57BL/6
Footpad Thickness (mm) (Mean+/- SEM)
Weeks post-primary infection
(1x10^4 L.m.-FV1 s.c./Footpad)
D
BALB/c-L.m.-FV1
BALB/c-L.m.-FV1_L.m.-FV1
Ear Lesion Area (mm^2) (Mean +/- SEM)
Days post-challenge
(2x10^5 L.m. i.d./Ear)
Parasite titer (-log/ear) (Mean+/-SD)
***

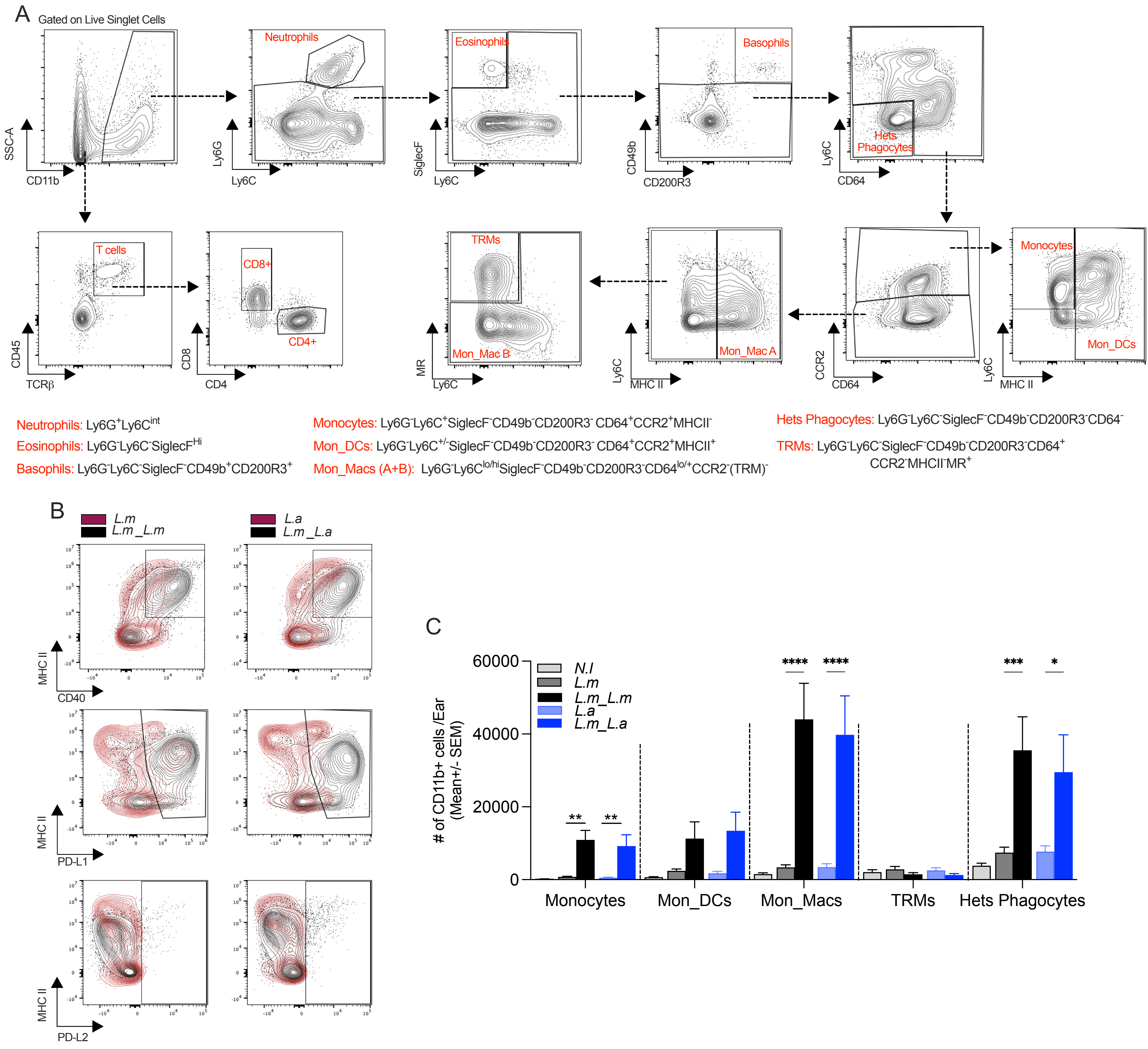
A
Gated on Live Singlet Cells
SSC-A
CD11b
Neutrophils
Ly6G
Ly6C
Eosinophils
SiglecF
Basophils
CD49b
CD200R3
Hets Phagocytes
CD64
T cells
CD45
TCRβ
CD8+
CD4+
CD8
CD4
TRMs
Mon_Mac B
MR
Mon_Mac A
MHC II
CCR2
Monocytes
Mon_DCs
Neutrophils: Ly6G+Ly6Cint
Eosinophils: Ly6G-Ly6C-SiglecFHi
Basophils: Ly6G-Ly6C-SiglecF-CD49b+CD200R3+
Monocytes: Ly6G-Ly6C+SiglecF-CD49b-CD200R3- CD64+CCR2+MHCII-
Mon_DCs: Ly6G-Ly6C+/-SiglecF-CD49b-CD200R3- CD64+CCR2+MHCII+
Mon_Macs (A+B): Ly6G-Ly6Clo/hiSiglecF-CD49b-CD200R3-CD64lo/+CCR2-(TRM)-
Hets Phagocytes: Ly6G-Ly6C-SiglecF-CD49b-CD200R3-CD64-
TRMs: Ly6G-Ly6C-SiglecF-CD49b-CD200R3-CD64+
CCR2-MHCII-MR+
B
L.m
L.m_L.m
L.a
L.m_L.a
MHC II
CD40
PD-L1
PD-L2
C
### of CD11b+ cells /Ear
(Mean+/- SEM)
60000
40000
20000
0
N.I
L.m
L.m_L.m
L.a
L.m_L.a
**
**
****
****
***
*
Monocytes
Mon_DCs
Mon_Macs
TRMs
Hets Phagocytes

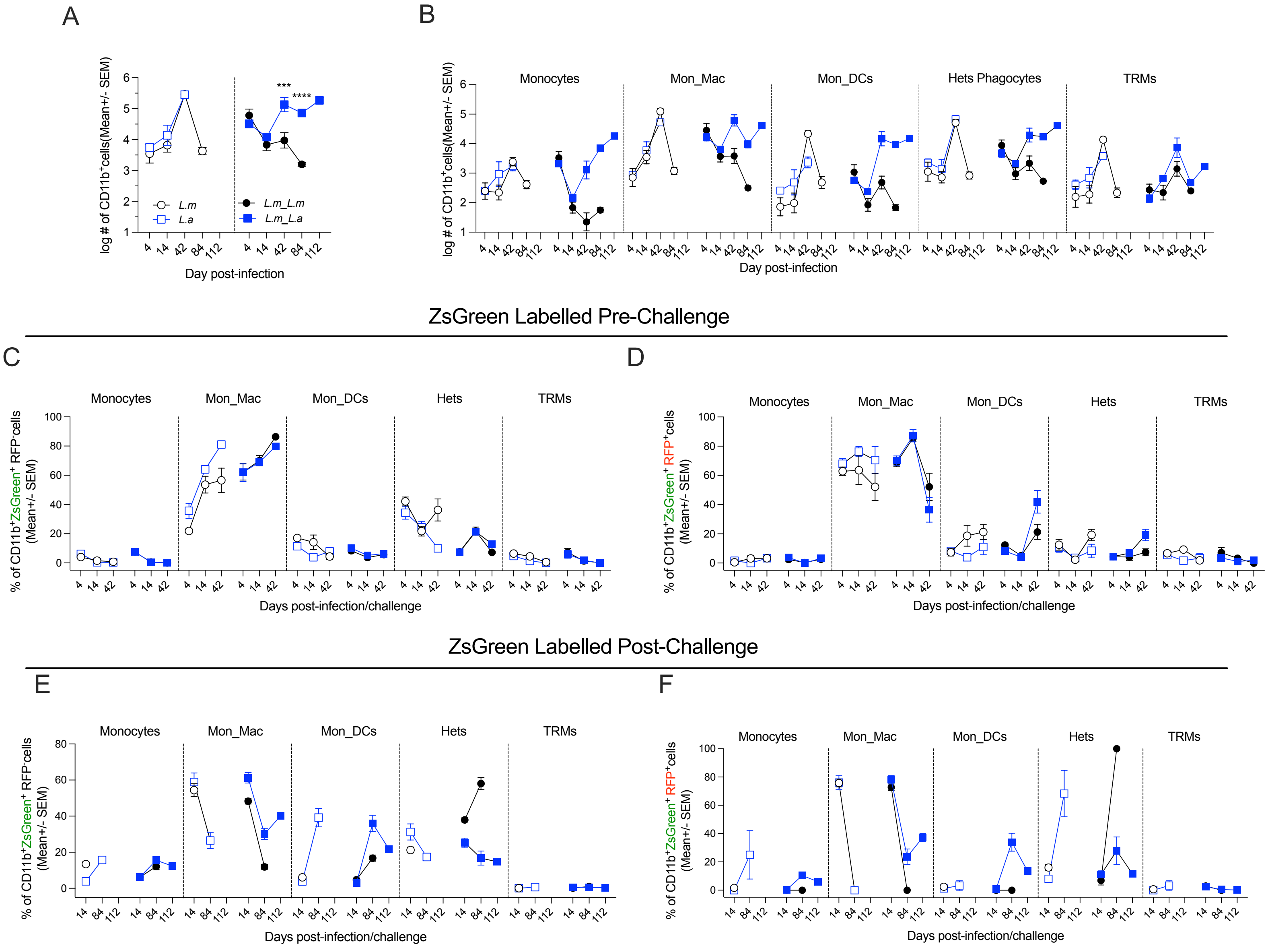
A
B
log # of CD11b⁺cells(Mean+/- SEM)
L.m
L.a
L.m_L.m
L.m_L.a
***
****
Day post-infection
Monocytes
Mon_Mac
Mon_DCs
Hets Phagocytes
TRMs
ZsGreen Labelled Pre-Challenge
C
D
% of CD11b⁺ZsGreen⁺ RFP⁻cells (Mean+/- SEM)
% of CD11b⁺ZsGreen⁺ RFP⁺cells (Mean+/- SEM)
Hets
Days post-infection/challenge
ZsGreen Labelled Post-Challenge
E
F

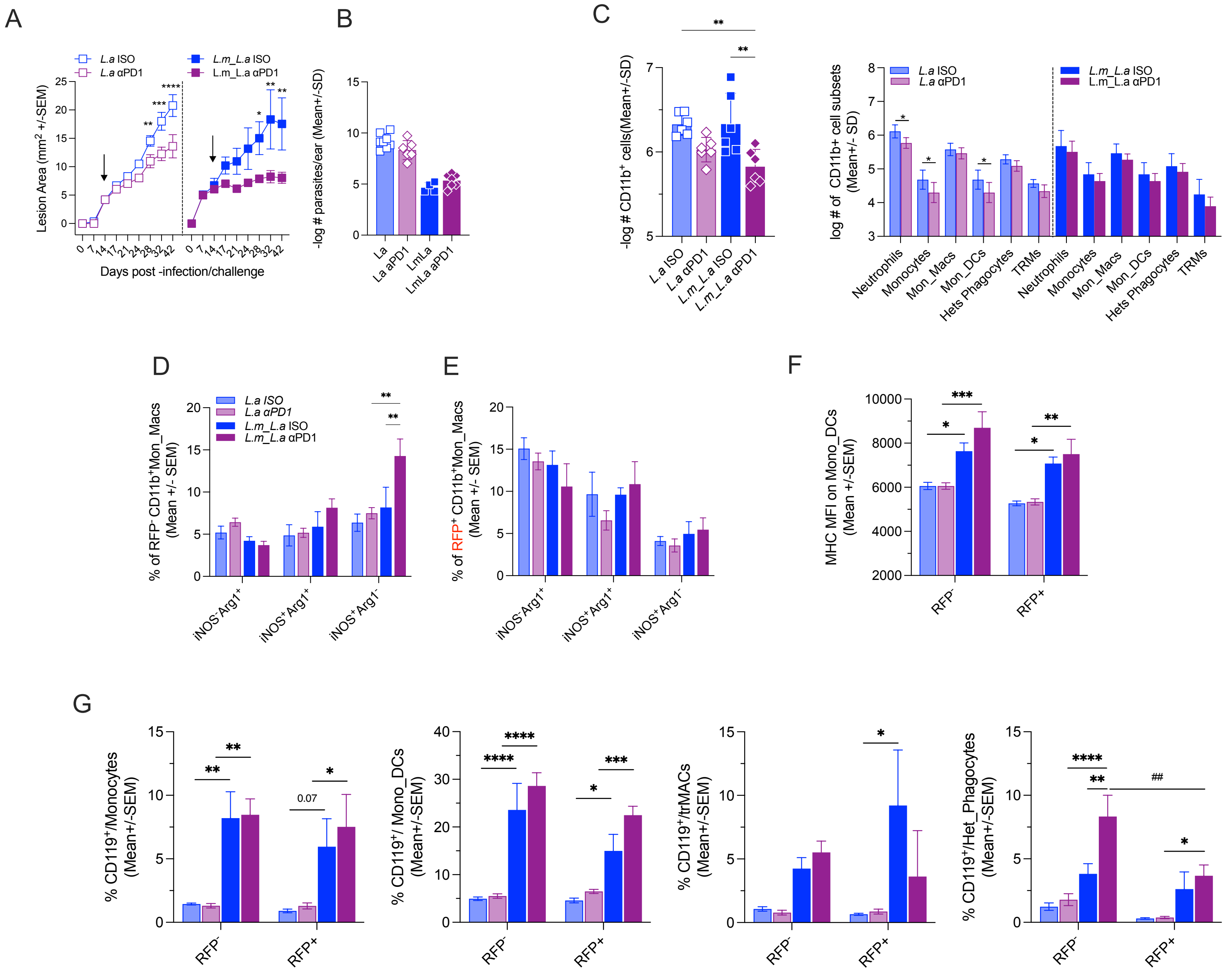

A
L.a ISO
L.a αPD1
L.m_L.a ISO
L.m_L.a αPD1
Lesion Area (mm² +/-SEM)
Days post -infection/challenge
B
-log # parasites/ear (Mean+/-SD)
La
La aPD1
LmLa
LmLa aPD1
C
-log # CD11b⁺ cells(Mean+/-SD)
log # of CD11b+ cell subsets (Mean+/- SD)
Neutrophils
Monocytes
Mon_Macs
Mon_DCs
Hets Phagocytes
TRMs
D
% of RFP⁻ CD11b⁺Mon_Macs (Mean +/- SEM)
iNOS⁻Arg1⁺
iNOS⁺Arg1⁺
iNOS⁺Arg1⁻
E
% of RFP⁺ CD11b⁺Mon_Macs (Mean +/- SEM)
F
MHC MFI on Mono_DCs (Mean +/-SEM)
RFP⁻
RFP+
G
% CD119⁺/Monocytes (Mean+/-SEM)
% CD119⁺/ Mono_DCs (Mean+/-SEM)
% CD119⁺/trMACs (Mean+/-SEM)
% CD119⁺/Het_Phagocytes (Mean+/-SEM)
0.07
##

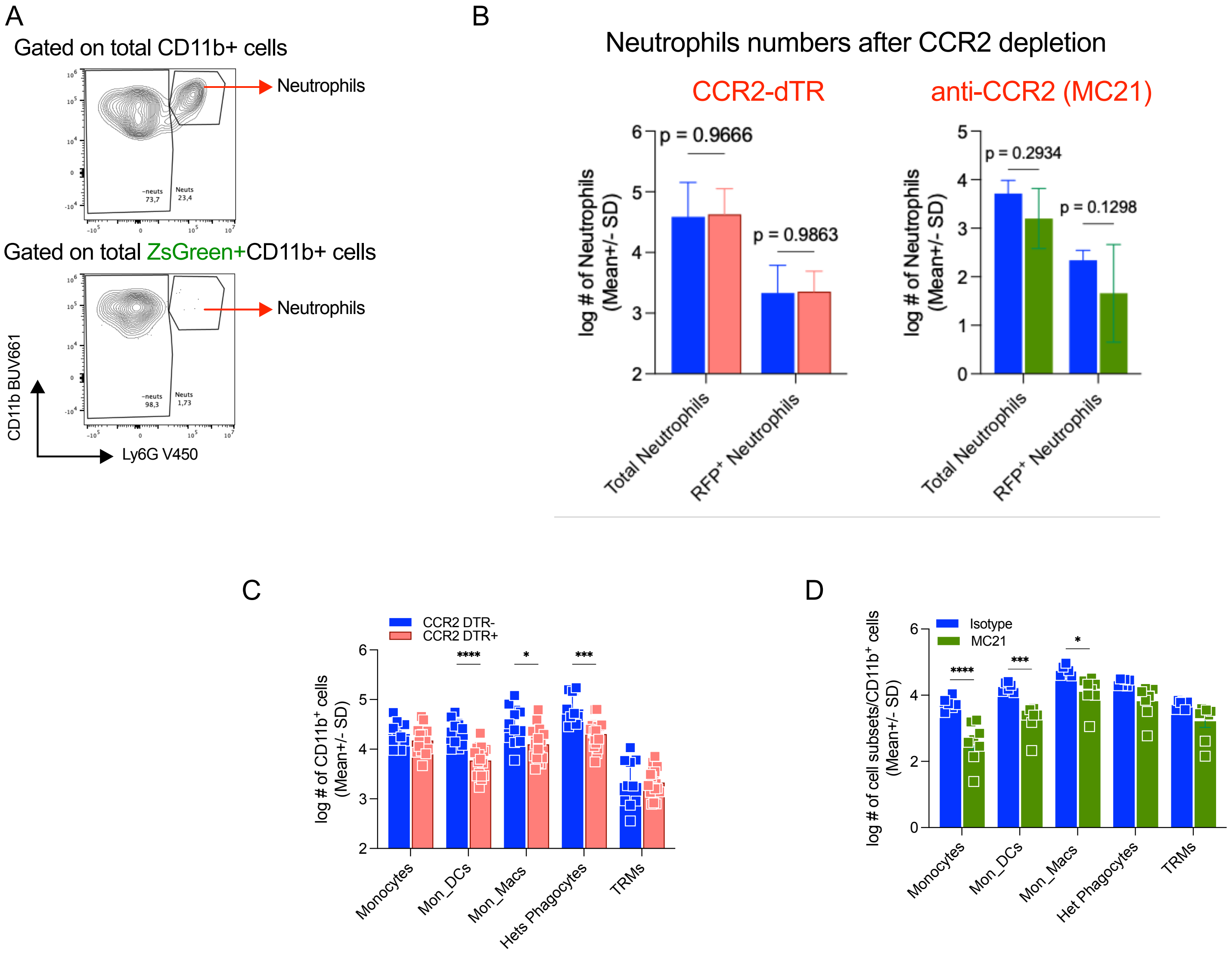

A
Gated on total CD11b+ cells
Neutrophils
Gated on total ZsGreen+CD11b+ cells
Neutrophils
CD11b BUV661
Ly6G V450
B
Neutrophils numbers after CCR2 depletion
CCR2-dTR
anti-CCR2 (MC21)
p = 0.9666
p = 0.9863
p = 0.2934
p = 0.1298
log # of Neutrophils (Mean+/- SD)
Total Neutrophils
RFP+ Neutrophils
C
CCR2 DTR-
CCR2 DTR+
log # of CD11b+ cells (Mean+/- SD)
Monocytes
Mon_DCs
Mon_Macs
Hets Phagocytes
TRMs
D
Isotype
MC21
log # of cell subsets/CD11b+ cells (Mean+/- SD)
Monocytes
Mon_DCs
Mon_Macs
Het Phagocytes
TRMs
